# In the same cell, the proteome defines cellular state and the transcriptome marks transitions

**DOI:** 10.64898/2026.08.25.746940

**Authors:** Marvin Thielert, Enes Ugur, Maximilian Zwiebel, Marc Oeller, Constantin Diekmann, Nils Eikmeier, Simon Suppinger, Lucas Diedrich, Sophia Steigerwald, Ankit Sinha, Prisca Liberali, Christoph Ziegenhain, Matthias Mann

**Author notes:** these authors contributed equally (Marvin Thielert, Enes Ugur, Maximilian Zwiebel, Marc Oeller, Constantin Diekmann).

## Abstract

Bulk transcriptome and proteome correlate only modestly, but this has not been investigated in the same cell or across cell-state changes. Here we introduce a scalable technology that quantifies thousands of proteins and transcripts in the same cell, separating RNA from protein by tip-based C18 capture and pairing full-length RNA sequencing with latest-generation mass spectrometry. In HeLa cells, transcript and protein abundances agree on the broad ranking within a cell (r = 0.45), but do not co-vary across the population (r = 0.038). In pluripotency transitions, only a third of matched transcripts and proteins change synchronously, yet the transcription factors defining each state stay tightly co-regulated. Transcript variance is several-fold larger than protein variance, reflecting transcriptional bursting and mRNA sampling noise. The proteome is thus the stable, low-noise definition of cell state, while the transcriptome marks cellular transitions; consequently, the proteome defines cell-state from far fewer cells.

**Highlights:**

- Multimodal workflow yields deep proteome and transcriptome from the same cells
- Transcript and protein rank similarly within a cell but are uncoupled across cells
- Only a third of RNA-protein pairs change together, yet state-defining TFs stay coupled
- Protein varies several-fold less than mRNA, defining state from far fewer cells

## Introduction

Single-cell transcriptomics has made the transcriptome the default readout of cellular state: cell types, developmental trajectories, and disease programs are now routinely defined from transcriptional profiles, often with the implicit assumption that RNA provides a complete representation of cellular function^1–3^. Yet proteins execute most cellular function, and transcript abundance predicts protein abundance only partially^4^: since the earliest genome-wide comparisons, mRNA has been shown to explain only a fraction of the variance in protein abundance, with the remainder shaped by translation rate, protein stability, and turnover^5–8^. This raises three questions: (i) how much of a cell’s functional, protein-level state is reflected in its transcriptome, (ii) how cell-to-cell variance is propagated from mRNAs to proteins during cellular state changes, and (iii) which of the two layers provides the more stable definition of cell state.

This decoupling is most consequential when cellular state is actively changing, as in the pluripotency continuum of embryonic stem cells (ESCs). The transition from naive through formative to primed pluripotency recapitulates successive stages of peri-implantation epiblast development and is accompanied by rapid changes in cellular signaling, metabolism, chromatin organization, lineage competence and transcription factor network^9–13^. This network is organized around pan-pluripotency stemness factors (OCT4, SOX2, SALL4 in mouse ESCs), together with changing stage-specific regulators, which diverge at the RNA and protein level^14–17^. Yet whether state-defining factors remain more tightly coupled between RNA and protein than the broader proteome - and how transcriptional changes are propagated, buffered, or delayed at the protein level - have only been inferred from bulk measurements and remain unresolved in individual cells.

Addressing the single-cell protein modality in depth has only recently become feasible. Single-cell proteomics has advanced rapidly through low-input sample preparation, more sensitive mass spectrometers, and data-independent acquisition^18–23^, pushing coverage from a few hundred to several thousand protein groups (protein forms that are distinguishable by mass spectrometry) per cell. The proteome can now be read at a depth approaching that of the transcriptome, yet it has remained a single-modality readout, reported in isolation rather than alongside the mRNA of the same cell.

Measuring both modalities in the same cell at high quality and scale, however, has remained technically challenging. Estimates of mRNA-protein concordance have almost all come from bulk measurements or inferred from single-cell datasets in which the two modalities are profiled in different cells and approximated computationally^24^. Single-cell strategies that combine modalities each involve a compromise: antibody-based readouts such as CITE-seq add only a few dozen pre-selected, mostly surface-localized epitopes and do not quantify the full proteome^25^; and physical splitting of single-cell lysates (nanoSPLITS)^26^ reaches genuine same-cell measurement at limited proteomic and transcriptomic depth and throughput. A recent preprint reports matched RNA and protein measurements from less than hundred cells, but at a depth and scale that cannot support systematic comparison across states^27^.

Here, we introduce a workflow that closes this gap by specific separation of target molecules rather than splitting of crude lysate. Briefly, cell lysis is followed by protein digestion, reverse transcription of mRNAs, amplification of cDNA and selective C_18_ capture of peptides, while the unbound cDNA flow- through is recovered for full-length transcriptome library construction. Pairing an adapted Smart-seq3 protocol with latest-generation mass spectrometry, we quantify deep transcriptomes and proteomes from the same individual cells. We apply this technology first to asynchronously growing HeLa cells to define a same-cell mRNA–protein correlatome, and then to mouse ESCs across the naive-to-primed transition to follow both modalities along a differentiation trajectory. Together, these systems allow us to investigate the unique information content of protein-level states and transcriptomes, and how cell-to-cell variance propagates between the two modalities.

## Results

### A single physical separation captures the deep proteome and transcriptome of the same cell

Reading both molecular modalities from the same cell has so far required a compromise. Because proteins cannot be amplified, existing same-cell approaches sacrifice one modality for the other: splitting the lysate halves the material available to each and limits depth^26^, whereas antibody-based readouts capture only a few dozen preselected epitopes rather than the full proteome. We instead separate the two molecular classes by their chemistry, so that neither is depleted (Figure 1A). This design rests on two developments. First, tryptic peptides bind a standard C18 solid phase tip (Evotip) while the untouched cDNA passes into the normally discarded flow-through, so that a single standard device performs the separation and recovers both modalities^28^. Second, we found that reverse-transcription (RT) and amplification of the single-cell RNA before tip-based separation was critical to achieve a deep single-cell transcriptome. Including RT and PCR in the same well after protease digestion and heat inactivation still yields full-length cDNA without compromising peptide recovery; this step was the most demanding to establish.

**Figure 1.**
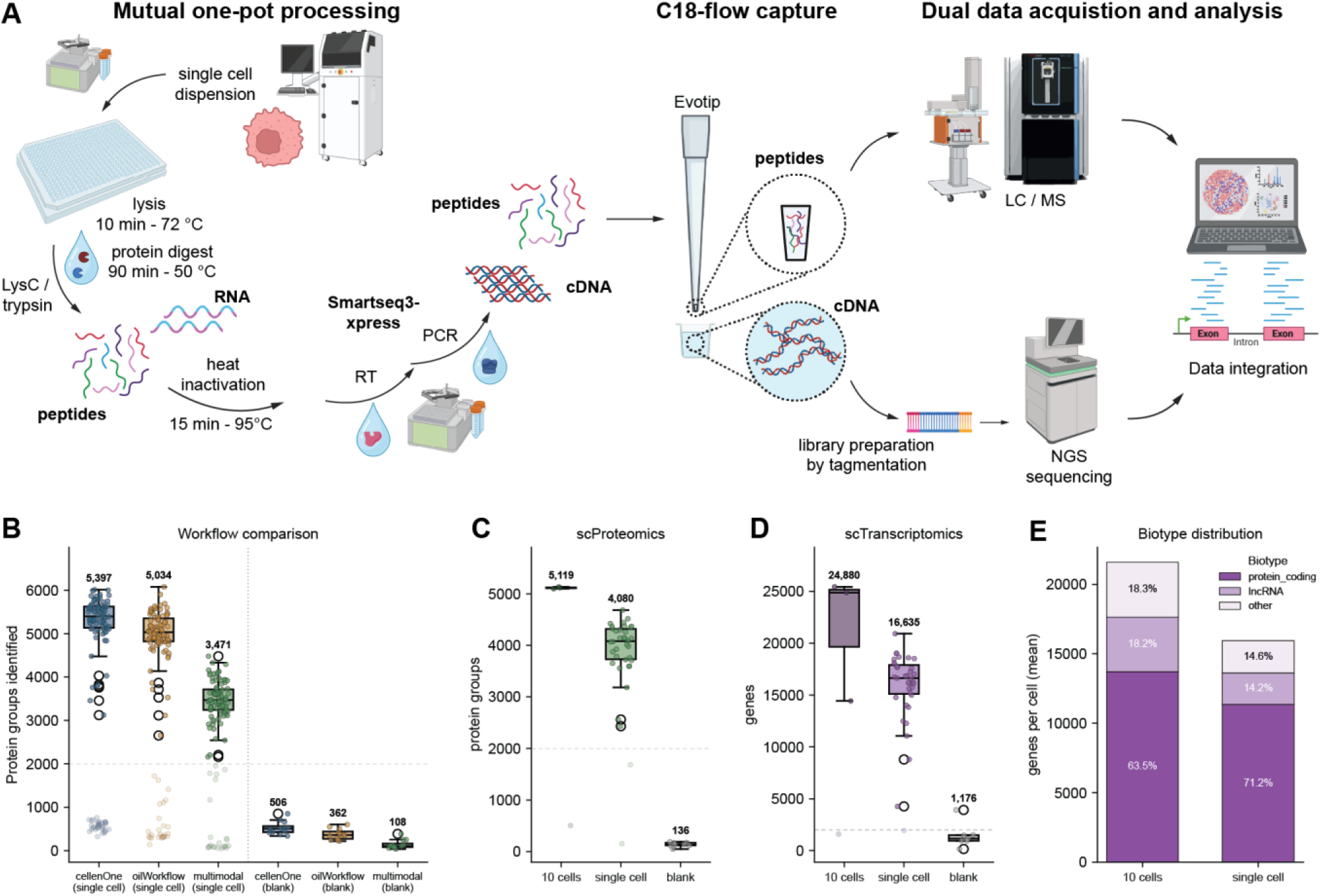
An integrated single-cell workflow quantifies the proteome and transcriptome of the same cell at scale and in depth across both modalities. (A) Schematic of the integrated single-cell proteomics and transcriptomics workflow. Single cells are dispensed (cellenONE) into oil-overlaid 384-well plates; a single combined lysis is followed by protein digestion and heat inactivation. Reverse transcription (RT) and pre-amplification (PCR) generate full-length cDNA. Tryptic peptides are captured on C18-tips (Evotip) while the unbound cDNA flow-through is recovered for tagmentation-based library preparation and NGS. Peptides are analyzed by LC-MS/MS on the Orbitrap Astral Zoom, and the two modalities are integrated computationally in our pipeline built on our Python-based AlphaPeptTools library. (B) Protein groups identified per single cell and blank control (blank) across the multimodal integrated workflow (multimodal) and single-cell proteomics, cellenONE and standalone oil workflow (n = 113 single cells and 12 blanks per condition). Boxes indicate median and IQR; the median is annotated. (C) Proteomic depth of the multimodal workflow per single cell, 10-cell equivalent, and blank (n = 36 single cells, 4 10-cell and 8 blanks per condition). (D) Genes detected in the multimodal workflow for the same samples and groups as in C. (E) Biotype composition of detected genes for 10-cell versus single-cell samples.

In the full workflow, individual cells were dispensed by either a piezo-acoustic dispenser (cellenONE) or by FACS into oil-overlaid 384-well plates and lysed (see Material and Methods). Lysis was followed by protein digestion, heat inactivation, RT and PCR before separating peptides and cDNA by C18 capture. Peptides were analyzed by latest-generation LC-MS/MS instrumentation, and cDNA flow-through was processed by tagmentation-based library preparation and sequencing. The paired measurements were bioinformatically integrated for each cell. Because neither modality is diluted or compromised, our design preserves the full cellular input for both readouts while retaining the throughput of a plate-based format. The order of steps was established by benchmarking alternative arrangements of digestion, reverse transcription (RT), pre-amplification (PCR), and C18 capture, while the sequence above yielded by far the best joint recovery (Figure S1A). Protein digestion before RT and PCR preserved the proteome recovery (Figure S1B-D), while transcriptome depth and library quality were retained when RT and PCR were done before C18 capture (Figure S1E, F). However, the cell sorter, sample preparation and back- end MS and NGS steps are independently swappable, making it a modular workflow that can be reconfigured according to the biological question. Paired measurements were integrated using a computational pipeline built on our in-house AlphaPeptTools framework, which imports single-cell modalities into AnnData^29^ and MuData^30^ objects within the scverse ecosystem^31^ (see Material and Methods).

Despite reading both modalities from one cell, the multimodal workflow even enhanced single-cell transcriptomics depth, presumably because protein digestion makes RNA more accessible, although we did not test this directly. It also preserved most of the proteomic depth of proteomics alone. In direct comparison with a cellenONE workflow^32,33^ and oil-overlay single-cell proteomics, we recovered a median of 3,471 protein groups per cell - roughly two-thirds of the depth of the standalone protocol, a modest cost for additionally recovering a full transcriptome from the same cell (Figure 1B). The identified protein groups were largely shared across the three workflows (91%, Figure S2A). Blank controls processed in parallel confirmed that identifications derive from genuine single-cell input rather than carry-over or background. Profiling the two modalities across single cells, 10-cell equivalents, and blanks in an independent experiment, the multimodal workflow reached a median of 4,080 protein groups (Figure 1C) and 16,635 genes per single cell (Figure 1D), rising to 5,119 protein groups and 24,880 genes in 10-cell samples while blanks remained in the range of a few hundred proteins. The single-cell proteome spanned about 4.6 orders of magnitude in abundance (Figure S2B), and a core set of 2,207 protein groups was detected in at least 90% of cells (Figure S2C). To our knowledge, this is the deepest single-cell proteome yet reported alongside a matched transcriptome, while its gene coverage far exceeds that of droplet-based single-cell RNA-seq methods, therefore both modalities reach great depth in the very same cell. Apart from the deep coverage, we capture full-length transcripts instead of just 3’ sequencing, enabling detection of splice junctions. The 10-cell pools were used only to benchmark the workflow and were excluded from all downstream single-cell analysis. Finally, the full-length RNA chemistry captured a broad range of transcript biotypes in the expected proportions^34^ (Figure 1E). Protein-coding messages dominated the detected genes (71% in single cells), with the remainder split between long non-coding RNAs (14%) and other non-coding classes (15%). At 10-cell input the protein-coding share fell to 64% as both non-coding classes rose (lncRNA 18%, other 18%), consistent with protein-coding detection approaching saturation at single-cell input so that additional depth recovers predominantly lower-abundant non-coding transcripts (Figure 1E). Read-mapping composition and gene-body coverage were consistent across single-cell and 10-cell samples and clearly separated from blanks (Figure S2D-I).

Together, these benchmarks establish a multimodal technology that quantifies thousands of proteins and transcripts from the same individual cell at a scale and depth suited for systematic RNA-protein comparisons.

### Single-cell matched profiling reveals weak mRNA-protein correlation per gene and lower protein variability

To establish a reference for matched RNA-protein analysis, we profiled asynchronously growing HeLa cells (n = 318 cells). Within each cell, comparing RNA and protein across the 3,904 genes detected in both modalities gave a modest positive correlation (median r = 0.45; Figure 2A, Figure S3B; metrics defined in Figure S3A). Measured within a single cell, this value recapitulates the modest concordance long reported between separate bulk datasets, confirming that the discordance is a property of the biology and not an artifact of comparing different samples.

**Figure 2.**
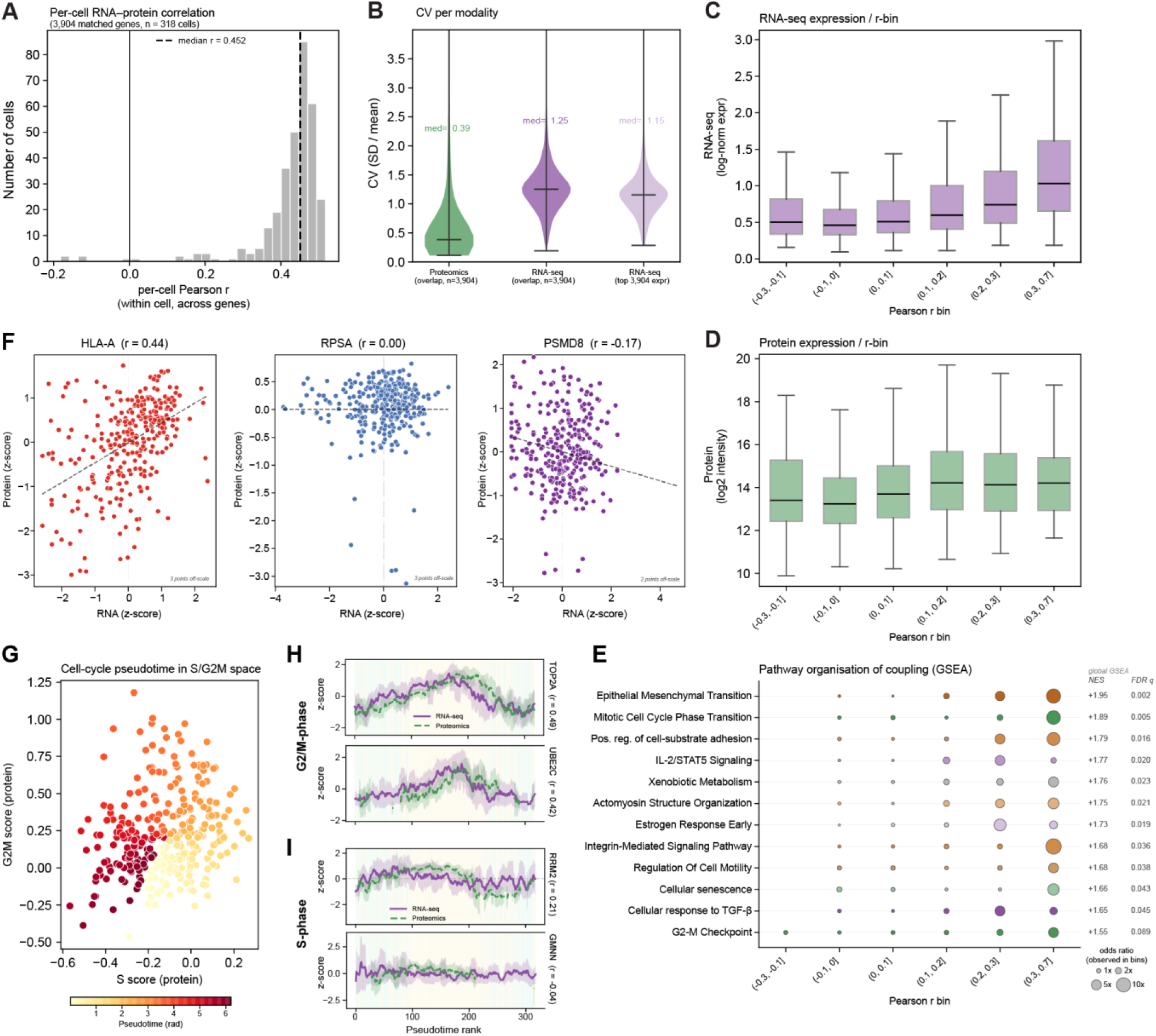
Matched single-cell profiling reveals the structure and limits of RNA-protein correlation across the cell cycle. (A) Distribution of per-cell RNA-protein Pearson correlations in matched HeLa single cells (n = 318 cells; 3,904 matched genes), each computed across genes. The four correlation metrics used in this study are defined schematically in Figure S3A. (B) Per-gene coefficient of variation (CV) by modality (y-axis clipped at the 98th percentile), showing markedly lower protein than transcript variability across cells. (C) RNA abundance (log-norm expression) levels stratified by per-gene Pearson r bin. (D) Same as C, but for protein abundance (log2 intensity). (E) Gene-set enrichment across Pearson r bins from C and D (dot size = −log10 FDR, color = gene ratio), identifying pathways whose members are consistently high- versus low-correlated. (F) Representative genes spanning the correlation range: strongly correlated HLA-A (left), non-correlated RPSA (middle), and anti-correlated PSMD8 (right), shown as per-cell z-scored protein versus RNA. (G) Cell-cycle trajectory reconstructed from protein-based S and G2/M scores colored by inferred pseudotime (rad) (n = 317 cells, 1 outlier removed). (H, I) Cell-cycle gene examples along pseudotime for G2/M markers (H) TOP2A (left) and UBE2C (right) and for S-phase markers (I) RRM2 (left) and GMNN (right). RNA and protein z-scores are visualized across the pseudotime rank from G.

The biologically informative comparison, however, runs along the other axis: for each gene, correlating its RNA and protein across the 318 cells. Computing the correlation for all matched transcript-protein pairs yields a distribution that is centered near zero (median r = 0.038, with a slight positive skew; Figure S3C and D). In other words, the abundance of a gene’s transcript is not predictive for the abundance of the corresponding protein product in an individual cell. Thus transcript and protein abundances agree on the broad ranking of gene products within a cell, but are uncorrelated with respect to cell-to-cell variation. Protein levels were markedly less variable across cells than transcript abundance (Figure 2B), indicating that the proteome is buffered against the cell-to-cell stochastic fluctuations that dominate the transcriptome. We had already reported lower protein than mRNA variability^23^, but these data establish this finding within the same cell, directly showing that the two modalities fluctuate independently. This buffering is expected on kinetic grounds: transcripts are made in short bursts and turn over within hours^35^, whereas proteins accumulate and persist far longer, so each protein level reflects a long time-average that damps out the transient, low-copy fluctuations of its mRNA.

Stratifying genes by their RNA-protein correlation bin showed that low correlation is not an artifact of low abundance, since transcript and protein levels spanned the full dynamic range within correlation bins, underlining the interpretation that this is a biological phenomenon not measurement noise. However, transcript abundance increased with correlation strength (Figure 2C), while protein abundance did not show any trend (Figure 2D). Gene-set enrichment across correlation bins showed that RNA–protein coupling is organized by pathway rather than distributed at random: members of the cell cycle (G2/M checkpoint, E2F targets, and mitotic spindle) were consistently well correlated, whereas weakly correlated genes were distributed more diffusely in oxidative phosphorylation and translation (Figure 2E). Representative genes spanning the range, from concordant (HLA-A) to uncorrelated (RPSA) to weakly anti-correlated (PSMD8), illustrate this per-cell relationship directly (Figure 2F). Together, these results establish a core same-cell correlatome and indicate protein-level buffering, rather than measurement noise, as the origin of the modest concordance.

### Correlation strength is pathway-dependent and pseudotime resolves phase-specific cell-cycle dynamics

The weak static correlation raised the question of whether coupled RNA-protein dynamics exist but are obscured when per-gene correlations are compared without temporal context. Because HeLa cells were unsynchronized, the population spanned the cell cycle, and we reasoned that ordering cells along this axis would expose dynamics invisible to static correlation. We reconstructed a cell-cycle trajectory directly from the protein modality, computing relative protein-based S and G2/M phase scores for each cell (marker sets from Tirosh et al.^36^) and assigning them a continuous cell-cycle pseudotime (Figure 2G, see Material and Methods).

Ordering matched measurements along this pseudotime recovered RNA-protein coupling that the static per-gene correlation had missed, and showed it to be phase- and gene-specific. Among canonical G2/M markers, TOP2A (r = 0.49) and UBE2C (r = 0.42) showed tightly coupled transcript and protein trajectories, with protein rising slightly behind the mRNA (Figure 2H). S-phase markers spanned a wider range: RRM2 (r = 0.21) showed comparable coupling earlier in pseudotime, whereas GMNN (r = −0.04) showed a flat transcript trajectory while its protein rose in S phase and decayed slowly thereafter (Figure 2I). Thus, the population-level enrichment of cell-cycle programs among highly correlated genes (Figure 2E) resolves, at single-cell and single-gene resolution, into a spectrum spanning from tight temporal coupling to fully independent transcriptional and translational regulation. Such decoupled trajectories may be a signature of post-transcriptional buffering and delayed protein turnover.

Together, these analyses show that the weak per-gene correlation in this proliferating population does not reflect an absence of coupling but rather its restriction to specific pathways and phases. This modality-specific temporal structure is invisible to single-modality measurements. The proteome both recovers the underlying trajectory and reveals where RNA and protein move together and where they diverge - establishing a steady-state baseline of RNA-protein coupling against which the dynamic pluripotency system is compared in the following sections.

### Matched single-cell profiling resolves the naive-to-primed transition in both proteome and transcriptome

We next applied our multimodal workflow to a dynamic system, profiling mouse embryonic stem cells (ESC) across the naive-to-primed pluripotency continuum at four different time points: naive (2i/LIF), a 6 h formative-induction intermediate (FGF2/ActA/XAV939), formative EpiLCs (2 days, 48 h)^12^, and primed EpiSCs (7 days, 168 h) (Figure 3A). Both modalities were recovered from every cell, with a median of 2,654 protein groups and 10,400 genes per cell across the 526 matched cells. Per-cell quality control confirmed high-quality datasets in both modalities across all four states, with low mitochondrial content and consistent feature (protein groups or genes) detection (Figure S4A–J). The number of identified features was slightly below the HeLa values as expected from these cells’ smaller size. Of features detected in at least 10% of a state’s cells, 3,627 genes were identified in both the transcriptome and the proteome (Figure S4K). The per-cell proteome and transcriptome depths were only modestly correlated (Pearson r = 0.39; Figure 3B). Since the cellenONE provides an estimated cell diameter, we related identified features directly to cell size: protein depth scaled more tightly with cell diameter (ρ = 0.73, n = 540) than the gene count did (ρ = 0.57, n = 612; Figure 3C, D), revealing that single cell size is a shared driver of both readouts and that the number of protein identifications is the more faithful single-cell size proxy. To resolve this scaling functionally, we next binned cells by diameter and observed that larger cells were enriched for genes related to mitotic/cell-cycle programs as well as proteins related to translation, ribosome-biogenesis and telomere maintenance (Figure S4L, M).

**Figure 3.**
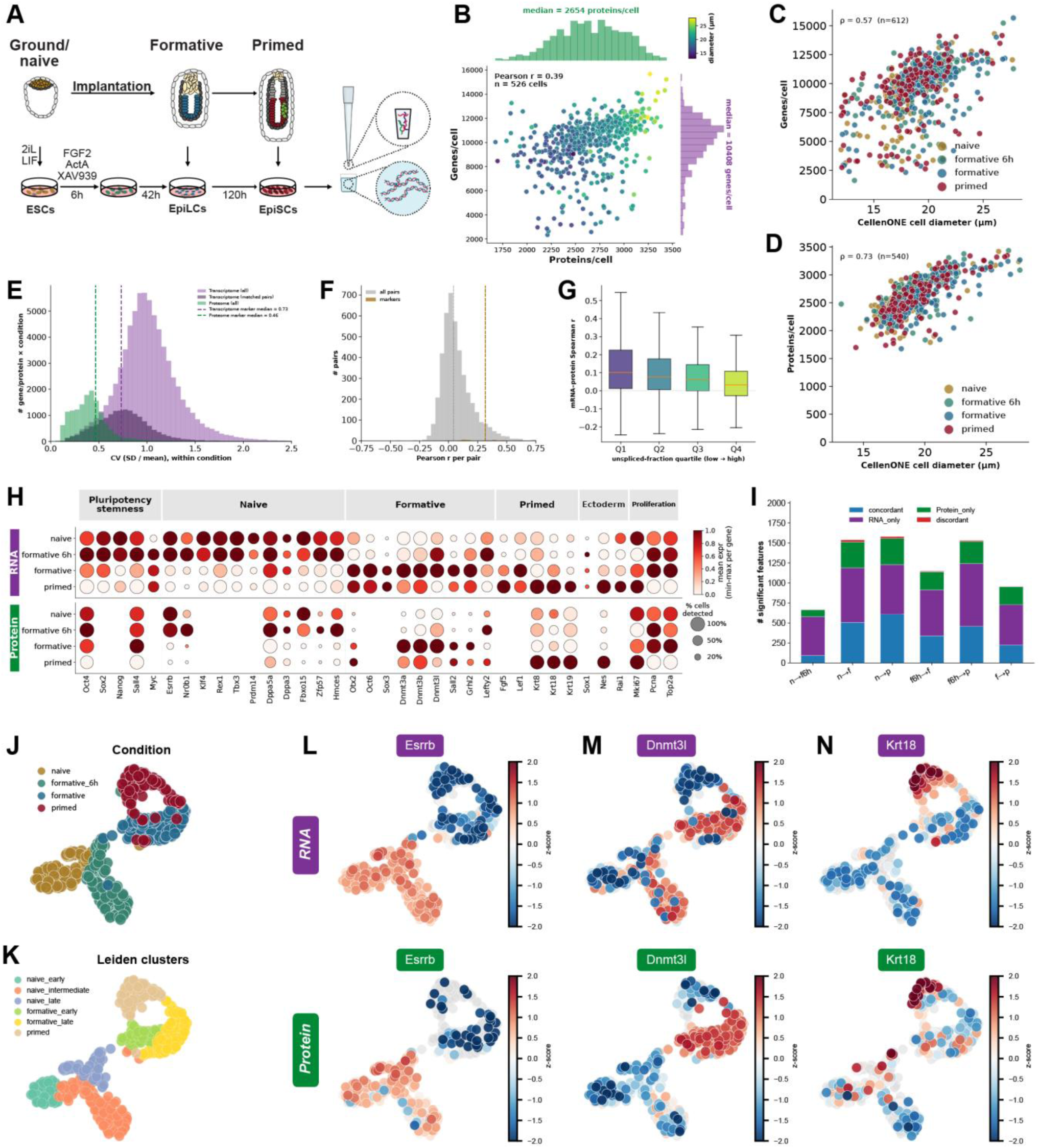
Matched multi-omic profiling resolves pluripotency state transitions and reveals tight marker coregulation despite overall limited RNA-protein correlation. (A) Schematic of the differentiation model: mouse embryonic stem cells (ESCs) progressing from the naive through formative (6 h, 48 h) to the primed state, with every cell profiled by the paired multimodal single-cell proteomics and transcriptomics workflow. (B) Per-cell transcriptome depth (genes detected) versus proteome depth (protein groups detected), colored by the cell diameter. (C) Genes detected per cell versus cell diameter and color-overlay for each timepoint. (D) Same as C, but for protein groups; the number of identified proteins tracks cell diameter more closely than gene count (Spearman ρ = 0.73 versus 0.57). (E) Per-feature coefficient of variation (CV) calculated within each state for the whole transcriptome (light purple), the overlapping transcript subset (dark purple), and the proteome; protein CV is 1.9-fold lower than matched transcript CV. (F) Distribution of per-gene RNA-protein Pearson correlations across matched single cells (n = 4,350 gene-protein pairs; median r = 0.044). Curated pluripotency markers (see also H) show higher per-gene correlation (n = 32 gene-protein pairs; median r = 0.32). (G) Per-gene RNA-protein Spearman correlation stratified by nascent (unspliced) transcript fraction (quartiles). (H) Curated lineage-marker dot plot across differentiation states (RNA top, protein bottom; dot size = fraction of cells detected, color = mean scaled expression). (I) Differentially expressed genes and protein changes across pairwise comparisons of pluripotency states (>1.4-fold change, FDR <0.05), split into shared and modality-exclusive changes. (J,K) Protein UMAP colored by differentiation state (J) and by protein Leiden cluster classification (K). (L–N) Per-cell z-scored expression of Esrrb (L), Dnmt3l (M), and Krt18 (N) on the protein UMAP coordinates (RNA top, protein bottom).

Three facets of the same kinetic cause distinguish the modalities. First, the protein-level buffering against transcriptomic variance seen in HeLa persisted in the differentiating population: quantified as the per-feature coefficient of variation (CV) across cells – our metric for cell-to-cell variability – the proteome was 2.5-fold less variable than the whole transcriptome within each condition (median CV 0.40 versus 0.99). When comparing only the matched proteome-transcriptome subset, the proteome was 1.9-fold less variable (median CV 0.40 vs 0.75) with identity defining markers showing an even smaller gap (1.6-fold lower, median marker CV 0.46 versus 0.73; Figure 3E).

Second, the per gene-protein correlation was gene-class specific: matched pairs for canonical stage markers were substantially better correlated than the transcriptome-wide background (Figure 3F), so the genes that define pluripotency state are precisely those whose protein tracks its mRNA. Ranking all pairs by their per-cell correlation and binning into deciles showed that coupling strength co-varied with abundance and pathway: both mean transcript and mean protein levels rose monotonically from the lowest to the highest decile in every state (Figure S5A, B), and gene-set enrichment along the same gradient placed ribosomal-subunit, actin-filament and secretory programs at the abundant, highly-correlated tail, while mitochondrial-gene-expression and RNA-processing sets concentrated at the weakly-correlated end (Figure S5C). The RNA-protein correlation weakened further as the unspliced (nascent) transcript fraction rose (Figure 3G). Because unspliced transcripts mark genes only recently induced, whose mature protein has not yet accumulated, this low correlation is expected as a snapshot of transcription that the proteome has not caught up to. Nascent transcripts anticorrelate with transcript abundance (r = -0.48), which contributes further to mRNA-protein decoupling due to additional transcriptional noise.

Third, despite being often offset in time, both modalities recapitulated the expected staging, resolving the ordered repression of naive factors (Esrrb, Nr0b1, Prdm14, Tbx3) and induction of formative and primed programs (Otx2, Dnmt3a/b/l, Fgf5, Lef1) in RNA and protein alike (Figure 3H). Yet differential testing across successive transitions exposed a pronounced asymmetry: the largest class of significantly changing features was detected in RNA alone (>1.4-fold, FDR <0.05, in pair-wise comparisons between successive states) throughout all differentiation time points, with similar concordant (protein and mRNA) and far fewer protein-only changes especially at 6h of differentiation. We rarely observed RNA-protein pairs with opposite enrichments between the differentiation time points (Figure 3I). Transcriptional change thus outpaces measurable protein change at every step. Lower variability and temporal lag are two faces of the same fact: protein is a time-averaged, buffered readout of a faster-fluctuating transcriptome.

To investigate biological reproducibility, we analyzed a larger single-cell proteomics (scp) only dataset (n = 884 cells) of the same E14 mESC line independently differentiated at three pluripotency stages (naive, formative and primed) but without the intermediate formative 6h state. Cell identities were resolved by stage-specific markers (Figure S6A) and after dataset integration they showed the same naive-to-primed transition (Figure S6B-F). Protein changes agreed in co-significance and direction (co- dir) in differential expression analysis (co-dir 92-96%, Figure S6G). This scp-only dataset provided slightly higher proteomic depth with a median of 3,569 versus 2,476 protein groups per cell, resulting in higher data completeness for curated markers and transcription factors (Figure S6H-J).

Projected into a UMAP of the protein data, the four differentiation stages occupied clearly separated regions ordered from naive through the two formative intermediates to primed, showing that the proteome alone resolves the continuum (Figure 3J). Leiden clustering of the protein-based nearest neighbors graph then resolved finer structure than the four nominal stages, splitting the naive cells into early, intermediate, and late sub-states and the formative cells into early and late states (Figure 3K). Overlaying individual markers on these coordinates, with RNA and protein displayed on the same cells (Figure 3L-N), showed that the two modalities demarcate the same identities but with a slight temporal offset. The naive transcription factor Esrrb declined in concordance on both modalities (Figure 3L), whereas Dnmt3l - the cofactor of the *de novo* DNA methyltransferases and an early formative marker - was already induced at the RNA level in the 6 h condition while its protein lagged behind (Figure 3M); Krt18 is a marker of the primed state but we clearly observed it on the protein level at the naive to formative transition as well (Figure 3N). An extended panel of eight lineage markers projected onto the RNA UMAP, with the protein-derived Leiden sub-states overlaid, confirmed that the protein-defined substructure maps coherently onto the transcriptional embedding: pluripotency factors (Oct4, Sall4) were high in the naive cluster and declined while Dnmt3b and Otx2 rose toward primed, concordantly in RNA and protein, although several markers were sparsely detected on protein level (e.g. Otx2 in 14% and Dppa3 in 23% of the cells, respectively) (Figure S5D). Overall, measuring both modalities therefore refines the boundaries of intermediate cell populations where the proteome captures a time shift of certain markers and likely also true cell identities, while the transcriptome covers more comprehensive cell identity defining markers.

### RNA and protein diverge in pathway dynamics along pseudotime and contribute to distinct axes of variation

Having established that the two modalities are often offset in time, we next asked which genes and programs drive the divergence. We first ordered cells along a continuous developmental axis independently in each modality. Diffusion pseudotime (DPT) provides an unsupervised ordering of cells by their respective diffusion-map distance from a naive root cell. The root cell is independent of the discrete pluripotency stage markers^37^. Both modalities were globally concordant, with RNA- and protein- derived DPT increasing together from naive to primed and diverging mainly across the early formative state, where protein lagged (Figure 4A).

**Figure 4.**
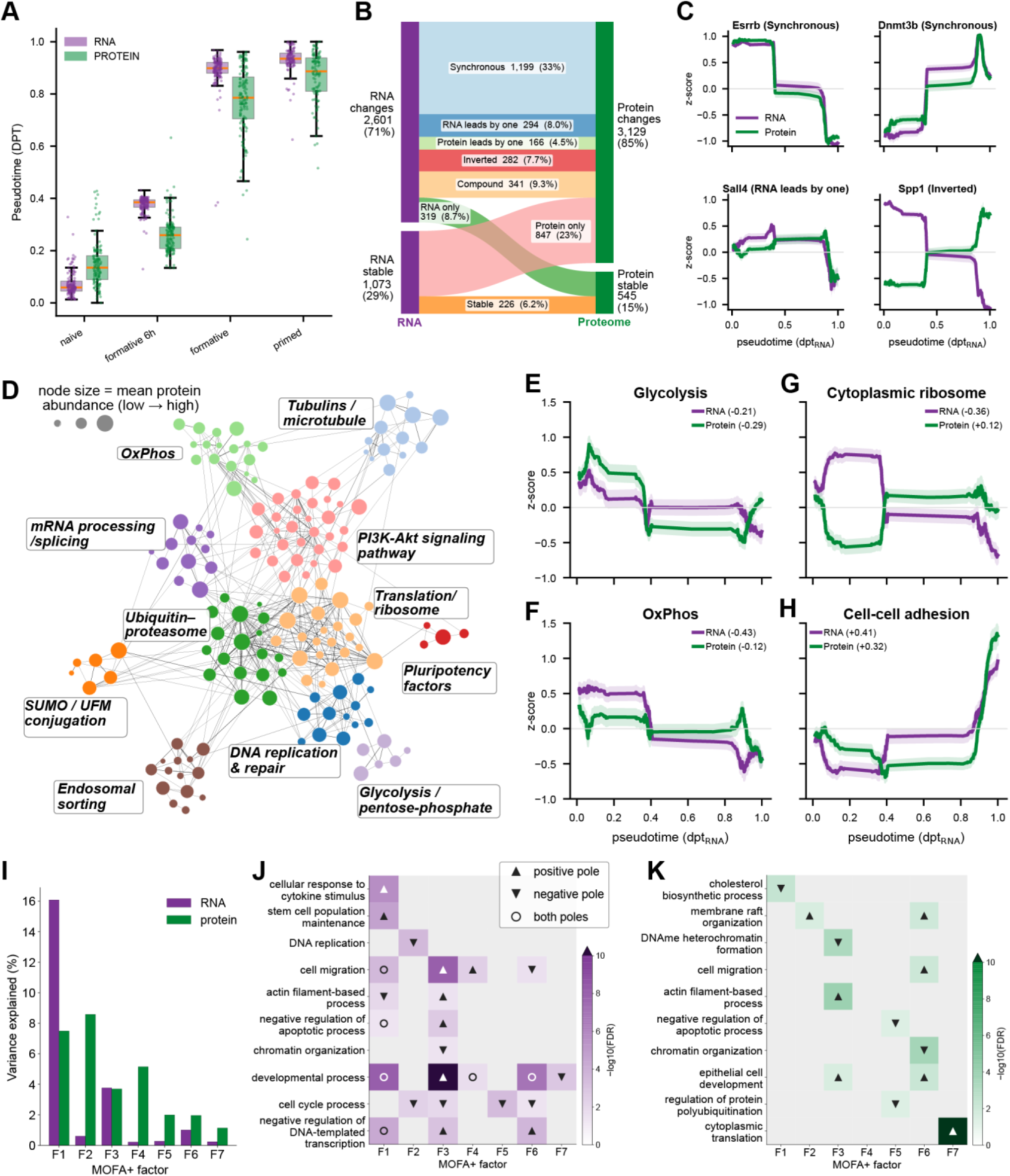
Joint pseudotime modeling resolves coupled and decoupled RNA-protein programs along differentiation. (A) Diffusion pseudotime per differentiation state for each modality (RNA purple, protein green); RNA and protein pseudotime are correlated (Pearson r = 0.94). (B) Per-gene classification of the RNA/protein trajectory patterns along five equal-count pseudotime bins (n = 3,674 matched genes), shown as the fraction of genes per class (synchronous, RNA-only, protein-only, RNA/protein leads by one, inverted, compound, stable). (C) Representative pluripotency/lineage trajectories - Esrrb and Dnmt3b (synchronous), Sall4 (RNA leads by one), and Spp1 (inverted) - as z-scored rolling mean ± SEM of RNA (purple) and protein (green) along RNA pseudotime. (D) STRING interaction network of the 155 gene-protein pairs whose RNA and protein pseudotime trajectories are decoupled (|Δρ| ≥ 0.40, FDR < 0.05, in total 201 gene-protein pairs of which 155 have at least three known interactions); nodes are colored by Louvain community and sized by mean protein abundance. (E–H) Functional program activity along pseudotime (z-scored per-cell score, rolling mean ± SEM) for RNA versus protein: glycolysis (E), oxidative phosphorylation (F), cytoplasmic ribosome (G), and cell–cell adhesion (H); the Spearman correlation with pseudotime is annotated for each modality. (I) MOFA+ variance explained per factor, split by modality. (J, K) GO:BP enrichment (-log10 FDR) of each MOFA factor’s top loadings for the transcriptome (J) and the proteome (K), as term × factor heatmaps (gray = not significant). Enrichment was run separately on each factor’s 50 most positively and 50 most negatively loaded features; markers denote the pole at which a term is enriched (upright triangle: positive, inverted triangle: negative, circle: both poles and therefore not directional).

Along this trajectory we classified each RNA-protein pair by how its two modalities changed across five equal-count pseudotime bins (n = 3,674 matched pairs; Figure 4B). One third of pairs changed synchronously (33%, 1,199 pairs) and only 6% were stable in both modalities, so most matched pairs changed in a modality-specific manner. Protein-only changes formed the largest of these classes (23%, 847 pairs), about 2.5-fold more frequent than RNA-only changes (9%). Of note, these results are based on significant changes across the pseudotime, unlike the previous pairwise comparison of each state with a defined fold change cutoff (|log2FC| > 0.5, FDR < 0.05; Figure 3I). Among pairs in which both modalities changed but with an offset, the transcript more often moves first (RNA led protein by one bin; for instance the RNA level increased already at the 6 h mark whereas the protein level followed only after 48 h) than the protein (8% versus 5%), and a further 8% were inverted, with RNA and protein moving in opposite directions. Representative trajectories illustrate the classes: Esrrb falling and Otx2 increasing synchronously in both modalities, Sall4 with RNA leading protein by one bin, and Spp1 inverted, with protein rising against falling RNA (Figure 4C).

We next asked whether the pairs where RNA or protein lags along the pseudotime - fall into coherent functional programs. In total we identified 201 shifted pairs of which 155 were known to have at least three interactors based on the STRING database^38^. After Louvain clustering, we resolved 11 functional communities, which revealed functional modules by GO enrichment, including pluripotency factors, translation/ribosome, oxidative phosphorylation, glycolysis, ubiquitin–proteasome, DNA replication, and cytoskeleton-related proteins (Figure 4D). The time shift of matched RNA-protein changes was therefore not scattered at random but concentrated in the machinery that sets protein levels after transcription - translation and ribosome biogenesis, and the ubiquitin-proteasome system - together with metabolism and chromatin remodeling; decoupling occurs precisely where protein output is set downstream of the transcript. Tracing the activity of these programs along pseudotime exposed modality-specific regulations: glycolysis declined in both modalities, oxidative phosphorylation-related RNAs decreased while corresponding proteins remained comparatively stable, and the cytoplasmic ribosome module moved in opposite directions in the two modalities across the transition (Figure 4E-H).

Finally, we decomposed the paired data with MOFA+, an unsupervised latent-variable model that generalizes factor analysis to multiple modalities^39^. MOFA+ breaks the data into a shared set of factors while measuring how much variance each one explains in the RNA and protein views separately (Figure 4I). Each modality was fit on its own features (2,000 variable transcripts; all detected proteins) and linked through the shared cells, where the decomposition recovered the same asymmetry. Of seven retained factors, factor 1 (F1) dominated and was shared, explaining the most variance in both views (16% RNA, 7.5% protein) and declining monotonically from naive to primed with concordant RNA and protein loadings (r ≈ 0.7-0.8); transcriptionally F1 was dominated by genes related to developmental program and at protein-level by metabolic reprogramming (Figure 4J, K). The remaining factors were strongly modality-skewed: beyond a shared primed-lineage axis, five factors explained variance almost exclusively in the proteome. Annotating the top loadings of these protein-specific factors by GO-term enrichment identified two programs prominent in the proteome yet essentially absent from the transcriptome - translation and ribosome biogenesis, chromatin organization and heterochromatin formation (Figure 4K). The translation/ribosome program is precisely the module our network analysis had flagged independently, in which RNA and protein moved in opposite directions along pseudotime (Figure 4E–H); the two independent analyses therefore converge on protein-level regulation of the translation machinery which the transcriptome does not report. Thus two independent analyses - the targeted network of decoupled pairs and the unsupervised factor decomposition - converge on the same conclusion: the proteome carries regulatory programs, centered on translation and chromatin, that the transcriptome does not report.

### The proteome defines cell state, the transcriptome maps the paths between states

We then asked how much the two modalities differ per gene-protein pair. As a measure of modality- specific variance we computed the squared CV (CV²)^8^. Per gene, transcript CV² exceeded protein CV² by a median of 4.9-fold (log₁₀[CV²_RNA/CV²_Protein] = +0.69), and by 2.8-fold for state-defining markers (Figure 5A). This lower per-feature protein variance suggests that fewer single-cell proteomes than transcriptomes are needed to observe a population’s heterogeneity. To test this, we truncated each modality’s PC space at 30% of explained variance, resulting in 6 components for the proteome and 26 for the transcriptome, and then asked how much of that component count is recovered from subsets of the cell population. The proteome recovered 80% of its full-sample count with 110 cells and was essentially complete at 300 cells, whereas the transcriptome required 345 cells for the same 80% and was still gaining components at 450 (Figure 5B). Recovering the same fraction of each modality’s component structure therefore takes substantially more cells for the transcriptome.

**Figure 5.**
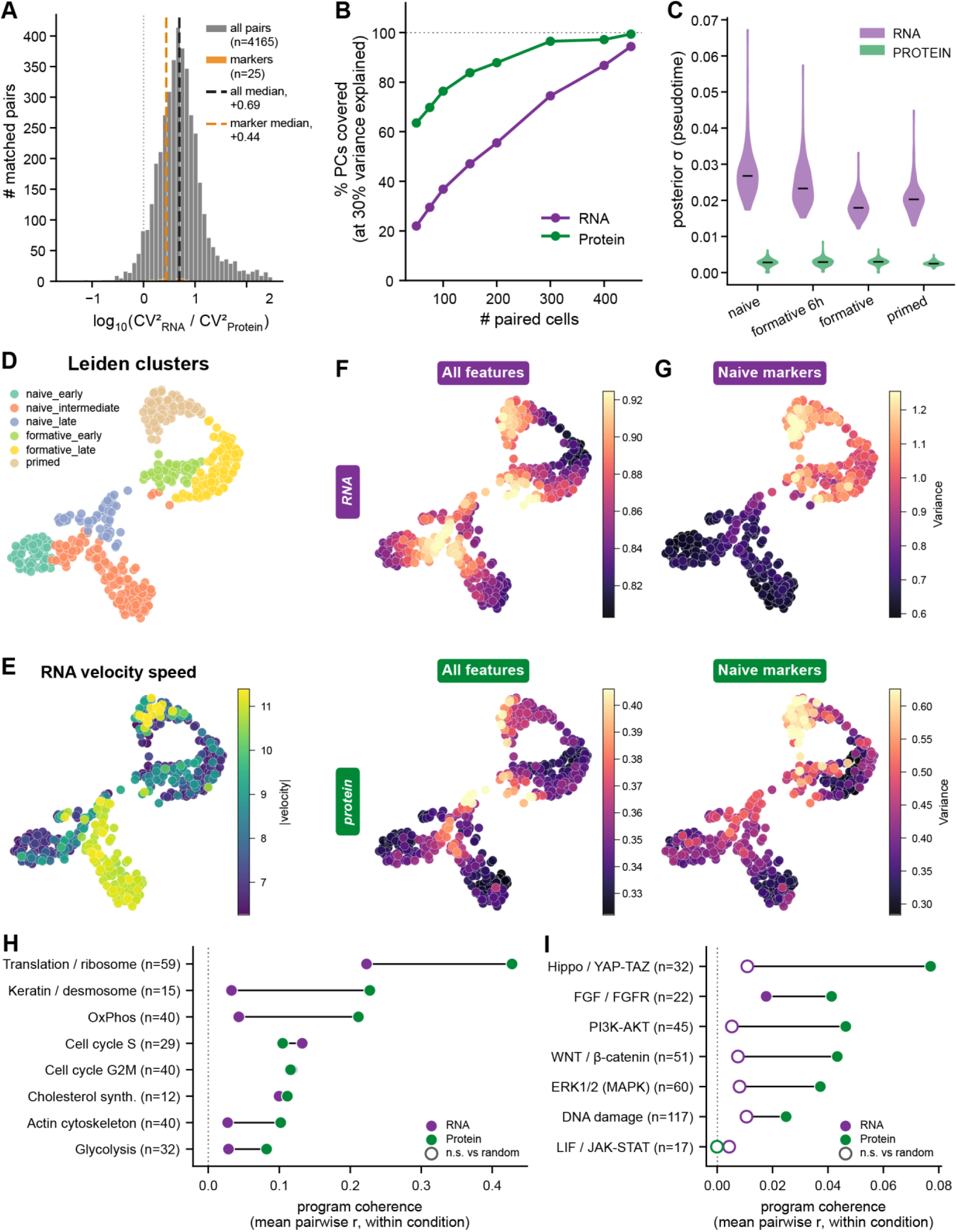
Greater transcriptome dynamics and proteome stability distinguish cellular potential from functional state. (A) Distribution of the per-pair log10 ratio of squared CV (CV²(RNA)/CV²(protein), computed within each state across unique matched gene-protein pairs (n = 4,165, gray); state-marker subset is overlaid (orange) and medians are marked (dashed lines). Transcript CV² exceeds protein CV² by a median of 4.9-fold across all pairs (log10 = +0.69) and 2.8-fold for state markers. (B) Percentage of each modality’s full-sample component count that is recovered from cell subsets, as a function of the number of paired cells sampled: the number of principal components explaining 30% of that modality’s variance at n cells, expressed as a percentage of the value at all 495 matched cells (mean of 20 subsamples; 1,742 matched gene–protein pairs with both members detected in ≥80% of cells). (C) Per-cell posterior σ of pseudotime position (Gaussian-process latent-variable model) per state and both modalities. (D) Protein UMAP colored by protein Leiden cluster. (E) RNA-velocity speed projected onto the protein UMAP. (F, G) Variance in UMAP space, that is the local CV (30-nearest-neighbor neighborhood) on the protein UMAP coordinates across all features (F) and across the naive-marker set (G) (RNA top, protein bottom). (H) Mean pairwise correlation among a cellular state program’s own features across cells, computed within each condition. High values mean the program’s members co-vary from cell to cell within a state, i.e. that modality reads a coherent axis of microstate variation. The same gene set is scored in both modalities (matched mRNA-protein pairs) and gene sets vary between n=12 and n=59. RNA purple, protein green; filled = above that point’s own size-matched random-set null, open = not significant. (I) same as in (H) for signaling related pathways (n varies from 17-117).

Our results also sharpen differentiation trajectory inference. A Bayesian Gaussian-process latent-variable model (GPLVM), which places each cell along a pseudotime axis as a probability distribution rather than a single point and reports the width of that distribution as the cell’s positional uncertainty, returned markedly tighter estimates from protein than from RNA (median posterior width 0.0036 versus 0.0212, in every condition; Figure 5C). The proteome thus locates a cell along the trajectory with far less ambiguity than the transcriptome.

Mapping cells onto the protein UMAP embedding, Leiden clustering resolved six substates spanning naive to primed (Figure 5D). Here the two modalities diverge in a characteristic manner. RNA velocity speed - the magnitude of the estimated per-cell rate of transcriptional change^40,41^, was elevated in the formative-entry (6 h) and primed compartments, marking these as the most transcriptionally dynamic (Figure 5E). We next overlaid per-cell, per-modality local variability on the same coordinates, computed as the coefficient of variation across each cell’s 30 nearest neighbors, either over all detected features of a modality (Figure 5F) or over the naive marker set alone (Figure 5G). Transcriptional variability peaked in the transitional compartments and protein variability, by contrast, remained uniformly lower and fell to a minimum at each population’s root (Figure 5F, G). Both measures describe the dispersion of individual features, and are therefore silent on how coherently a whole functional program is read out.

Besides cell identity, we further asked whether both modalities differently resolve cellular state, the relative abundance of functional programs. To test this, we computed for each functional program the co-variation of its protein-mRNA matched members from cell to cell within a single differentiation state. This measures whether a modality resolves cellular state once the differentiation axis itself is removed. We assessed significance against size-matched random sets of matched pairs, whose coherence is near zero and equal in the two modalities (0.008 for RNA, 0.015 for protein). The proteome was the more coherent readout for six of eight programs, most strongly for translation and ribosome biogenesis (0.43 versus 0.22), the keratin–desmosome module (0.23 versus 0.03) and oxidative phosphorylation (0.21 versus 0.04), whereas both cell-cycle programs were read at least as coherently from RNA (S phase 0.13 versus 0.11; G2/M 0.12 versus 0.12) (Figure 5H). Applying the same measure to signaling pathways (Figure 5I) showed that coherence was five- to ten-fold lower than for the metabolic programs; the transcriptome was indistinguishable from random for six of seven pathways, and although the proteome remained significant for six of seven (Hippo/YAP-TAZ 0.077, PI3K–AKT 0.046, WNT/β-catenin 0.043, FGF/FGFR 0.041, ERK1/2 0.037, DNA-damage response 0.025), it did so at uniformly small values, showing that cellular signaling state is still more coherently resolved by proteins. Together, the proteome therefore reports where a cell is, with low uncertainty and sharp boundaries between states, whereas the transcriptome’s variance and velocity concentrate precisely where cells are changing - encoding not the settled state but the direction and potential of movement between states. In the same cell, then, the proteome defines cellular state while the transcriptome maps the possible paths between them.

## Discussion

Single-cell transcriptomics has been so productive that it is now natural to treat the cell as a “bag of RNA”^42^, an abstraction in which mRNA is taken to suffice for describing cell state and identity. Our same-cell measurements challenge that view. Read from the very same cell, the transcriptome and proteome are only weakly correlated and, per gene, essentially uncorrelated (Figure S3C and D). The proteome varies far less and defines cell state more stably than mRNA. The transcriptome is a stochastic, anticipatory signal of the changes a cell is making, while the proteome is the low-noise readout of the functional state it is in. This highlights the need for true multi-modal measurements to gain a detailed understanding of cellular states and trajectories.

Our technology combines three properties not previously available together - same-cell measurement of the proteome and transcriptome, analytical depth in both, and plate-based throughput - by physically separating the two molecular classes rather than splitting the lysate, so that neither is depleted to obtain the other. From a single cell it quantifies a median of 4,080 protein groups and 16,635 read-aligned transcripts, to our knowledge the deepest matched single-cell proteome reported to date.

Read from the same cell, the two modalities stand in a defined quantitative relationship. Across genes within a cell they are only modestly correlated (r = 0.45), close to the concordance long inferred from separate bulk datasets^4,43^; per gene across cells the correlation is essentially zero, yet the proteome varies markedly less than the transcriptome^23^. These are not separate observations but two faces of the same kinetics: protein is a time-averaged, buffered result of transcriptional bursting^44–46^, so its cell-to-cell fluctuations are damped, and its abundance reflects a cell’s accumulated history rather than its instantaneous transcriptional state. Ordering cells along a protein-derived cell-cycle pseudotime makes this concrete, resolving the flat static correlation into gene- and phase-specific trajectories in which protein trails, tracks, or diverges from its mRNA - structure invisible to either modality alone.

Our same-cell measurements resolve a long-standing ambiguity regarding a predictive kinetic principle for when the transcriptome can substitute for the proteome. Because transcripts are produced in bursts and turn over within hours^6,47,48^, whereas proteins turn over roughly five-fold more slowly, with a median half-life on the order of tens of hours^6,49^, the protein level can represent a time-averaged integral of its mRNA. This single kinetic relationship predicts which genes agree between the two modalities and which diverge, without invoking gene-specific regulation. The modalities agree for abundant, stably expressed genes measured across cells - the identity-defining transcription factors whose protein tracks its mRNA closely in both HeLa cells and the pluripotency continuum. They diverge, predictably, in three regimes: for low-copy, bursty transcripts, where sampling noise dominates the RNA layer but is escaped by the far more abundant proteome; for recently induced genes, where the nascent (unspliced) transcript fraction is high but protein has not yet accumulated; and for genes whose output is set after transcription - the translation and ribosome machinery, protein turnover, and metabolic reprogramming that our network and factor analyses independently identify. It follows that the transcriptome is a faithful proxy for cell state for the abundant, constitutively expressed genes, and an unreliable proxy where post-transcriptional regulation predominates. Weak mRNA-protein correlation is therefore not measurement noise or an artifact of bulk averaging, but the expected signature of a buffered, time-delayed system, here measured directly in the same cell.

Thus transcriptomic atlases, and the cell identity and state definitions built upon them, report the anticipatory layer of gene expression: they capture the direction of a cell’s change with high sensitivity but define its present state with several-fold greater noise, requiring approximately 3-fold more cells to resolve the same population structure. The proteome provides the complementary measurement, a stable, low-noise definition of the present state with sharper boundaries between substates where even a single different cell can be informative and reliably discriminated^50^. The two measurements are complementary and cannot be substituted for one another; the temporal offset between them is itself informative. Because both are measured from the same cell, this dataset also provides the reference needed to calibrate the RNA-to-protein prediction models on which transcriptome-only atlases implicitly depend, and a template for extending same-cell measurement to additional molecular layers. Determining cell state from protein and trajectory from RNA within the same cell therefore constitutes a general approach for studying how cells commit to and execute fate decisions.

Extending the technology to a dynamic system, we profiled mouse embryonic stem cells (ESCs) across the naive-to-primed pluripotency continuum. The buffering seen in HeLa was not a property of a homogeneous state: within every differentiation stage, protein abundance remained markedly less variable than the matched transcripts, while state-defining transcription factors were well correlated for both modalities, indicating a more efficient post-transcriptional and translational control^51,52^. Despite the overall low correlation, the protein embedding resolved transition states equally well compared to the transcriptome with a slight time shift. Consistent with the proteome scaling with cell content, the number of proteins quantified per cell tracked cell diameter more tightly than gene count did. That protein sharpens rather than only reflects transcriptional state likely follows from its lower cell-to-cell noise rather than from the proteome carrying more information per cell. The modest mRNA-protein correlation long documented in bulk^4–6^ or on unmatched single-cell efforts^24^ thus persists and is now directly quantified.

Ordering the paired cells along a diffusion pseudotime trajectory revealed that coupled flow from transcript to protein is the exception: only 33% of matched pairs changed synchronously, while most changed in one modality alone, and where both changed with an offset the transcript more often moved first. This decoupling was not uniform but concentrated in defined machinery such as the translation/ribosome apparatus, oxidative and glycolytic metabolism, and the ubiquitin-proteasome system. An unsupervised factor decomposition^39,53^ recovered that both modalities resolved a shared developmental axis alongside protein-driven regulatory networks related to protein turnover as well as overall chromatin organization and specifically heterochromatin formation. This is in line with our previous finding of a global increase of heterochromatin-driving proteins in primed pluripotency^17^. Our observations also support the central role of translational control in pluripotency, where global protein synthesis is held low despite high ribosome biogenesis and ribosome remodeling gates fate transitions^15,54,55^. The RNA-protein time gap itself can be largely accounted for by protein turnover^56^ and by post-transcriptional attenuation of complex subunits^4,57^. The classical naive-to-primed metabolic switch^58^ appears here as transcript-protein decoupling rather than a coupled shift - a distinction that transcript abundance alone cannot resolve, as it does not report flux.

Yet the two modalities are not merely offset in time; they agree most precisely where it matters most for identity. Against the near-zero gene-wide correlation, the transcription factors that define each state remain tightly correlated between mRNA and protein. This reconciles their two roles: the transcriptome can chart the paths a cell may follow because the regulators that specify where those paths lead are read faithfully at both modalities, anchoring RNA’s forward-looking signal to the proteome’s definition of the present state. Decoupling is therefore selective - it spares the state-defining core while relaxing across the bulk of the proteome, whose levels are set after transcription.

Treating each modality as a measurement of cell state, we found that transcript cell-to-cell variability exceeded protein by 4.9-fold on the squared-CV scale (Figure 5A), and the proteome required substantially fewer cells to cover the heterogeneity of a population than the transcriptome (Figure 5B). As a consequence, population heterogeneity on the transcript level is much more broadly reflected and cell state is more definitive based on the proteome modality (Figures 5 H and I). Transcriptional variability, by contrast, was largest in the transition compartments, consistent with the proteome acting as a temporal low-pass filter on a burstier transcriptome. This spread reflects two effects: the sampling (shot) noise inherent to low transcript copy numbers, which the far more abundant proteome largely escapes, and buffering of mRNA fluctuations by translation and degradation. This within-cell result directly quantifies a principle - that transcriptional bursts are buffered before they reach a stable proteome^45,59,60^, a measure which single-modality measurements could only suggest^8^. With future technological advances, proteomics holds the potential to extend the analysis to protein isoforms and in principle the same-cell design can be extended to further molecular layers. Importantly, the proteome depth reported here will likely improve further as single-cell proteomic sensitivity and throughput continue to advance rapidly. We therefore anticipate for the near future that our same-cell technology will enable us to quantify low-abundant regulators which are missed at the current depth.

Our central claim should be read narrowly: the proteome provides the more stable definition of cell state, not that the transcriptome is dispensable. RNA remains the more accessible, higher-throughput and genome-wide layer, indispensable for unbiased discovery and for capturing the anticipatory dynamics that precede protein change; the “bag of RNA” abstraction and the atlases built on it are complemented, not overturned, by a protein view. Indeed, the two modalities converge on the same population structure: cells order almost identically along a pseudotime derived from either layer (Spearman ρ = 0.81) and the protein-derived Leiden clusters map coherently onto the RNA embedding. Single-cell proteomics therefore offers an alternative view of the same latent cellular state. That a single cell proteome might suffice to define its state is an expectation from the low noise of the proteome, not yet a demonstration. Several technical limitations will further improve in the future. The proteome depth still falls short of bulk proteomics and of the deepest single-cell methods reported. Therefore low-abundant regulators, including many signaling kinases and transcription factors, may be incompletely sampled. The throughput, although substantially higher than prior same-cell proteomics methods, remains orders of magnitude below droplet-based scRNA-seq. This is partly offset by the greater per-cell stability of the proteome, which requires far fewer cells to recover the same biological insights (Figure 5B). Because cells must be dissociated and viable at the point of sorting, the technology does not yet extend to fixed tissue or spatially resolved samples. Finally, missing values in the transcriptome and proteome require imputation and batch correction for downstream analysis; this missingness is in part a property of low-input single-cell measurement rather than a purely technical artifact, and its treatment influences quantitative comparisons between modalities. Because both modalities are read from the same cell, the dataset also provides a benchmark for methods that model missing values and cross-modal prediction directly, and a foundation for asking how far a single cell’s proteome alone can define its state and anticipate its fate.

## Acknowledgments

We thank all Mann Labs members for insightful discussions. We thank Wolfgang Enard for fruitful discussions and his highly valuable input on single-cell RNA sequencing technologies. This work is funded by the Max Planck Society for the Advancement of Science, by the Bavarian State Ministry of Health and Care through the research project DigiMed Bayern (www.digimed-bayern.de); the Swedish Research Council (2022-01471, C.Z.), the Frontier Grant of the Department of Medical Biochemistry and Biophysics (Karolinska Institutet, C.Z.); and the Swiss National Science Foundation (SNSF, grant number 223263, P.L.). M.O. was supported by the HORIZON-MSCA-2023-PF-01-01 project HETLEWY, no. 101151819. Some of the graphical illustrations used in the manuscript were prepared using BioRender.com.

## Author contributions

M.T., E.U. and M.M. conceived the project. M.T., E.U., M.Z., M.O. and C.D. developed the multimodal workflow, designed and carried out experiments. C.D. and M.Z. performed sequencing. E.U., Si.Su., and P.L. conceptualized the biological system. M.T. and So.St. conceptualized the single-cell proteomics data acquisition. M.T., E.U., M.Z., M.O., C.D., N.E., L.D. and A.S. performed data analysis. M.T., M.O. and C.D. curated data. M.T., E.U., C.Z., and M.M. supervised the project. M.T., E.U. and M.M. wrote the manuscript, and all authors reviewed the manuscript.

## Declaration of generative AI and AI-assisted technologies in the manuscript preparation process

During the preparation of this work, the authors used Claude Opus 4.5 and 5 by Anthropic to improve the readability and conciseness of the manuscript. After using this tool/service, the authors reviewed and edited the output and content as needed and take full responsibility for the content of the published article.

## Resource availability

### Lead contact

Further information and requests for resources and reagents should be directed to and will be fulfilled by the lead contact, Matthias Mann.

### Materials availability

This study did not generate new unique reagents.

## Data and Code Availability

### Data Availability

All data will be made available upon publication of the manuscript.

### Code Availability

All code presented herein as part of the multimodal workflow will be available upon publication of the manuscript on GitHub under the permissive Apache license.

## Material and Methods

### HeLa cell culture

Human epithelial carcinoma cells (HeLa, ATCC S3 subclone) were cultured in DMEM supplemented with 2 mM L-glutamine, 10% FBS, and 1% penicillin-streptomycin. Cells were grown to about 80% confluency and harvested with Accutase (Invitrogen) for 1 min at 37 °C. For single-cell sorting, cells were washed three times with ice-cold PBS, passed through a Flowmi cell strainer (70 μm mesh size), and diluted to a final concentration of 200 cells/μL in ice-cold PBS. Defined cell numbers were taken from these stock suspensions for single-cell and bulk proteomics experiments.

### Murine ES cell culture

Naive E14 mouse embryonic stem cells (mESCs) were cultured in serum-free medium consisting of N2B27 [50% Neurobasal medium (Life Technologies), 50% DMEM/F12 (Life Technologies)] supplemented with 2i [1 μM PD0325901 and 3 μM CHIR99021 (Axon Medchem, Netherlands)], 1000 U/ml recombinant leukemia inhibitory factor (LIF; Millipore), 0.3% BSA (Gibco), 2 mM L-glutamine (Life Technologies), 0.1 mM β-mercaptoethanol (Life Technologies), N2 supplement (Life Technologies), B27 serum-free supplement (Life Technologies), and penicillin-streptomycin (100 U/ml and 100 μg/ml, respectively; Sigma). To resolve the transition from naive pluripotency through the formative to the primed state, cells were sampled at four time points along a single differentiation time course: 0 h (naive mESCs), 6 h (a formative-induction intermediate state), 2 days (corresponding to formative EpiLCs), and 7 days (corresponding to primed EpiSCs). Differentiation was induced by switching naive mESCs into serum-free N2B27 medium lacking 2i, LIF, and BSA, supplemented with 10 ng/ml Fgf2 (RCD Systems), 20 ng/ml Activin A (RCD Systems), 5 μM XAV939 (Sigma) and 0.1x Knockout Serum Replacement (KSR; Life Technologies) (differentiation medium). The 0 h sample was collected directly from naive culture conditions 2-3 passages after thawing a respective ESC stock. The 6 h samples were harvested from cells maintained in differentiation medium without intervening medium change. EpiLCs (2 days) were harvested after one medium change at 24 h, and EpiSCs (7 days) were harvested after medium changes at 24 h, 3 and 5 days. All pluripotent stem cell models were cultured on 0.2% gelatin-treated 6-well plates. For cell sorting, cells were harvested, washed three times with ice-cold PBS, passed through a Flowmi cell strainer (70 μm mesh size), and diluted to a final concentration of 200 cells/μL in ice-cold PBS, matching the procedure executed for HeLa cells. Cells were in general not cultured longer than 10 passages and were tested for mycoplasma contamination.

### Single-cell proteomics cellenONE sample preparation

400 nL lysis buffer (100 mM triethylammonium bicarbonate buffer (TEAB), 0.2% DDM, 10 ng / μL LysC (Wako), 20 ng / μL trypsin platinum (Promega)) was dispensed in each well of a 384-well twin.tec PCR plate (Eppendorf) using a Mantis (Formulatrix) liquid handler. Individual cells were sorted into the lysis buffer using a cellenONE microfluidic dispenser (Cellenion). After sorting, the PDC was cleaned by sciCLEAN and three PDC flush steps to avoid contamination by remaining cell debris during the subsequent incubation. Cells were lysed and digested at 50 °C for 90 min inside the cellenONE. During incubation each well was rehydrated by the cleaned PDC with 500 drops in cycles of about 5 minutes. The interval between cycles was empirically optimized to avoid both drying of the well and excessive accumulation of volume over the incubation period. After digestion, the plate was cooled to 4 °C and acidified with 5 μL 0.1% formic acid in water. Peptides were loaded onto Evotips and stored at 4 °C until LC-MS measurement.

### Combined single-cell proteomics and transcriptomics sample preparation

Single cells were processed with a multimodal workflow that recovers tryptic peptides and full-length cDNA from the same cell, combining a miniaturized oil-based proteomics preparation with the Smart- seq3xpress chemistry (protocols.io, https://www.protocols.io/view/smart-seq3xpress-yxmvmk1yng3p/v3)^61^. All sub-μL dispensing was performed in 384-well twin.tec PCR plates (Eppendorf) using a Mantis (Formulatrix) liquid handler. Reagent compositions and per-well volumes are provided in Supplemental Table S1. The 10-cell and 20-cell samples used for benchmarking were processed identically to single cells, except that the cellenONE dispensed 10 or 20 cells into a single well instead of one. For blank controls, no cell was dispensed; all subsequent processing was identical to the single-cell workflow.

Plates were prepared by first dispensing 2 μL of hexadecane (Sigma) per well as an evaporation barrier, followed by 0.2 μL of lysis master mix per well, yielding final in-well concentrations of 0.20% n-dodecyl- β-D-maltoside (DDM), 0.005 U/μL SEQURNA RNase inhibitor (Lucigen), 0.438 μM biotinylated oligo- dT30VN primer (412), and 1.75 mM dNTPs (each) in nuclease-free water. Plates were briefly centrifuged and then cooled to freeze the oil. Individual cells were sorted into the lysis buffer/oil overlay using a cellenONE microfluidic dispenser (Cellenion). Plates were spun down briefly and then frozen and stored at −80 °C.

For processing, plates were thawed during centrifugation, and incubated at 72 °C for 10 min to lyse cells and release proteins and RNA. 0.3 μL of digestion master mix was then dispensed per well, bringing the final composition to 25 mM Tris-HCl (pH 8.3), 0.20% DDM, 8 ng trypsin platinum (Promega), and 4 ng LysC (Wako) per well. Protein digestion was carried out for 90 min at 50 °C, followed by heat inactivation at 95 °C for 15 min and a brief centrifugation. Reverse transcription (RT) was performed by addition of 0.2 μL of RT mix per well, giving final concentrations of 25 mM Tris-HCl (pH 8.3), 30 mM NaCl, 1.0 mM GTP, 2.5 mM MgCl₂, 1.5 mM DTT, 0.75 μM template-switching oligo (TSO; M462), and 2 U/μL Maxima H Minus reverse transcriptase (Thermo Fisher Scientific), and initial incubation at 42 °C for 90 min, 10 cycles at 50 °C and 42 °C for 2 min each, and 85 °C for 5 min for enzyme inactivation. cDNA was pre-amplified by addition of 0.8 μL of PCR mix containing 1x SeqAmp PCR buffer, 0.025 U/μL SeqAmp polymerase (Takara), and 0.5 μM each of forward (441) and reverse PCR primers (414), with initial incubation at 95 °C for 1 min, followed by 10-20 cycles of 98 °C for 10 s, 65 °C for 30 s, and 68 °C for 4 min, and a final extension at 72 °C for 10 min. The number of pre-amplification cycles was empirically optimized for each cell type and recorded per plate. If not stated otherwise, 18 cycles were used. After cDNA pre-amplification, the combined peptide/cDNA reaction was acidified by addition of 7 μL of 0.1% formic acid (FA) per well. Tryptic peptides were captured on Evotips Pure (Evosep) following the manufacturer’s loading protocol using the Bravo robot for automatization^62^; each well was subsequently rinsed with 8 μL of Evosep Buffer A and the rinse loaded onto the corresponding Evotip as well. The Evotip flow-through (∼16-17 μL per well), containing the pre-amplified cDNA, was collected in a fresh sealed 96-well plate (twin.tec LoBind, Eppendorf). Loaded Evotips were stored at 4 °C until LC-MS/MS analysis (see below), while cDNA-containing plates were shipped on dry ice to the laboratory of Christoph Ziegenhain (Karolinska Institutet, Stockholm, Sweden) for the remaining Smart-seq3xpress library construction steps - initial clean-up, tagmentation, indexing PCR, pooling, and final clean-up - with reaction volumes scaled to the cDNA pre-amplification volume as detailed in Supplemental Table S1 (see *Transcriptomics library preparation from flow through*).

Oligonucleotide sequences (Integrated DNA Technologies) used in this study were:

- Oligo-dT30VN (412): /5Biosg/ACGAGCATCAGCAGCATACGA-T(30)-VN (RNase-free HPLC)
- TSO (M462): /5BiosG/AGAGACAGATTGCGCAATG-NNNNNNNN-WW-rGrGrG (RNase-free HPLC)
- Forward PCR primer (441): TCGTCGGCAGCGTCAGATGTGTATAAGAGACAGATTGCGCAA*T*G (HPLC)
- Reverse PCR primer (414): ACGAGCATCAGCAGCATAC*G*A (HPLC)

### Transcriptomics library preparation from flow through

Smart-seq3xpress library construction steps were adjusted from previous protocol^61^ and performed on the Evotip flow-through: initial clean-up, tagmentation, indexing PCR, pooling, and final clean-up. Reaction volumes scaled to the cDNA pre-amplification volume are outlined in Supplemental Table S1. In detail, plates containing the Evotips flow-through with cDNA were cleaned up using 0.6x MGIEasy Clean beads (MGI) and eluted in 6 µL. 2 µL clean-up elution were transferred into a new plate and 1 µL 1x tagmentation buffer (10 mM Tris-HCl pH 7.5, 5 mM MgCl2, 5% DMF, 0.005 µL per well TDE1 Tn5 (Illumina)) was added. Plates were pulse-spinned to 1000 x g and incubated at 55 °C for 10 minutes. After tagmentation, 0.5 µL 0.28% SDS was added. Subsequently, 5 µL of custom Nextera index primers (0.2 µM, see Supplemental Table S2) carrying 10-bp dual indexes were added to each well followed by the addition of 4 µL PCR Mastermix (1x Phusion Buffer (Thermo Fisher Scientific), 0.01 U/µL Phusion HF Polymerase (Thermo Fisher Scientific), 0.2 mM dNTP each and 0.025 % Tween-20 (Sigma)). Plates were incubated for 3 minutes at 72 °C, 30 seconds at 98 °C, 18 cycles (10 seconds at 98 °C, 30 seconds at 55 °C, 30-60 seconds at 72 °C), and 5 minutes at 72 °C in a thermal cycler. Samples were pooled by spinning out each plate into a 300 mL robotic reservoir (Nalgene) using a custom 3D-printed scaffold (pulse-spin to about 200 x g). The pooled libraries were purified with home-made 22% PEG at 0.75x. Library quality was assessed by the Qubit dsDNA HS Kit and Tapestation D5000 Kit.

### cDNA library sequencing

Libraries were sequenced on MGI DNBSEQ G400RS platform (version 1.8.2.2293 software). Single-stranded circularized DNA libraries were created following the MGIEasy Universal Library Conversion Kit (MGI). 80 fmol of the resulting library were used for DNA nanoball (DNB) making (reagents provided in sequencing kit) immediately before sequencing.

### MS data acquisition on the Orbitrap Astral

Peptides were analyzed on an Orbitrap Astral and Orbitrap Astral Zoom mass spectrometer (Thermo Fisher Scientific) coupled to an Evosep One liquid chromatography system (Evosep Biosystems). Loaded Evotips were eluted using the Whisper Zoom 80SPD method (16.3 min active gradient) and peptides separated on an Aurora Elite analytical column (5 cm × 75 μm i.d., 1.7 μm C18 particles; IonOpticks) maintained at 50 °C in a column oven. The LC system was interfaced to the mass spectrometer via an EASY-Spray ion source equipped with a FAIMS Pro interface (Thermo Fisher Scientific), with the FAIMS operated at a single compensation voltage of −40 V and a carrier gas flow of 3.5 L/min. Electrospray ionization was performed at 1,900 V with the RF lens set to 40.

Data was acquired in data-independent acquisition (DIA) mode. MS1 survey scans were recorded in the Orbitrap analyzer from 380 to 980 m/z at a resolving power of 240,000 (at m/z 200), with a normalized AGC target of 500% and a maximum injection time of 100 ms. MS2 scans were acquired in the Astral analyzer across 150-2,000 m/z, covering the 380–980 m/z precursor range with 45 variable-width isolation windows optimized on the basis of precursor density using py_diAID^63^. Each MS2 scan used a maximum injection time of 26 ms and a normalized AGC target of 800%, and isolated precursors were fragmented by higher-energy collisional dissociation (HCD) at a normalized collision energy of 25%.

### Proteomics raw data analysis with DIA-NN

Raw mass spectrometry files were converted to mzML format and searched with DIA-NN v1.8.1 or v2.5.0^64^ against an in silico predicted spectral library generated from the Mus musculus reference proteome (UniProt UP000000589, downloaded January 17, 2024; 54,858 protein groups in 94,914 protein isoforms, 6,466,515 precursors) or Human reference proteome (UniProt UP000005640, downloaded March 2022; 100,727 protein groups in 172,613 protein isoforms, 6,254,585 precursors). The library was reannotated against the FASTA database during the search. Mass tolerances were fixed at 4 ppm for MS1 and 6 ppm for MS2, with a scan window radius of 7. Match-between-runs was enabled, allowing identifications to be propagated across all single-cell runs based on retention time and precursor mass alignment. Precursor and protein group identifications were filtered at 1% FDR, and quantification was performed using fixed-width peak centers with interference removal from fragment elution curves disabled. Protein grouping was performed in heuristic mode, and retention-time profiling was enabled. Protein, gene, and precursor quantities were exported as cross-run matrices for downstream analysis. Reported blank numbers were searched together with single-cell samples and single-cell samples were searched without carrier runs. Carrier runs of 10 cell and 20 cell were only used for number comparisons.

### Sequencing data processing

Raw FASTQ files were processed using the zUMIs pipeline^65^ (version 2.9.7 or newer) with default settings. Reads were filtered for low-quality barcodes and UMIs (reads with more than 5 bases < phred 20 were discarded), and UMI-containing reads were parsed by detection of the Smart-seq3 tag sequence (ATTGCGCAATG), allowing up to two mismatches. Within zUMIs, reads were mapped to the human (hg38, GRCh38) or mouse (mm39, GRCm39) reference genome using STAR (version 2.7.3). Genes were assigned using Ensembl annotations for human (GRCh38, release 95; GENCODE v29) and for mouse (GRCm39, release 110; GENCODE vM33). Error-corrected UMI counts were calculated per gene, and zUMIs reported for both, read-aligned and UMI-deduplicated gene-count matrices. Transcript counts in this study are reported on read-aligned counts unless stated otherwise zUMIs were also used to downsample cells to equal raw sequencing depth to facilitate method benchmarking.

### Single-cell multi-omics data analyses and integration

Downstream analysis was performed in Python using the alphapepttools package (v0.2.0) on top of the AnnData/scanpy ecosystem; the full notebook and configuration files are available at GitHub (see *Code availability*). Briefly, the zUMIs gene-level count matrices for scTranscriptomics and the DIA-NN protein-group matrices for scProteomics were imported into an AnnData object, and per-cell metadata (experiment, plate, 384-well position, quadrant, Evotip position, and group class) were parsed from proteomics raw file names. Raw intensities were retained as a separate layer for each modality, while for all subsequent analyses a log2-transformed layer was generated for proteomics and a log1p- transformed layer for transcriptomics after counts were normalized to 10,000 per cell.

Quality control was performed at both the cell and the feature level on both modalities proteomics and transcriptomics. Cells were flagged as outliers when their log-transformed median intensity or fraction of detected protein groups deviated by more than three median absolute deviations from the cohort median, following the recommendations of Heumos, Schaar et al.^66^, and outlier cells were excluded from further analysis.

Proteome preprocessing: Further, protein groups were filtered by completeness, retaining only features detected in at least 10% of cells. Missing values in the filtered matrix were imputed on features passing the completeness filter using a random-forest-based imputation^67^ (MissForest-style iterative imputation using extremely randomized trees; parameters: n_estimators=100, max_iteration=10) if not stated otherwise. Plate-to-plate technical variation was corrected using pyComBat with plate as the batch covariate. The contribution of known technical and biological covariates to global variance was quantified before and after batch correction by principal-component regression on the top 50 components^68^, confirming that batch correction reduced plate-associated variance without collapsing biological signal. Finally, PCA, k-nearest-neighbor graph construction, and UMAP embedding were performed using scanpy (v1.12.2) defaults on the batch-corrected, imputed matrix.

Transcriptome preprocessing: Ensembl gene identifiers were mapped to gene symbols and restricted to protein-coding, long non-coding, transcribed-pseudogene and mitochondrial biotypes, and counts sharing a symbol were summed. Cells were filtered to have at least 1,000 detected genes, at least 5,000 total counts, and no more than 20% mitochondrial reads. The 3,000 most highly variable genes were selected, scaled (clipped at 10 s.d.), and used for PCA analysis (50 components); a neighborhood graph (5 neighbors, 30 principal components), a UMAP embedding (mind_dist = 0.8), and Leiden clustering were then computed. Plate-associated batch effects were corrected with Harmony (harmonypy v0.2.0)^69^ on the principal components using plate as the batch covariate, and the neighborhood graph and UMAP were recomputed on the Harmony-corrected embedding. Marker overlays are drawn from the unimputed layers, z-scored across detected cells only, with undetected cells shown in grey.

Cross-modal integration: The processed proteome and transcriptome AnnData objects were combined into a single MuData (mudata v0.3.9) container. Because proteins and transcripts were acquired from the same physically isolated cells, the two modalities were paired by their shared unique single-cell identity of the plate and the 384-well source coordinate (plate_384-well) - yielding truly paired cells with both proteome and transcriptome measurements; cells measured in only one modality were retained as modality-specific observations. CellenONE index-sorting parameters were also matched to the unique single-cell identifier. To enable feature-level comparison across modalities, transcript symbols were matched to protein groups through UniProt: gene symbols (including recorded synonyms) were mapped to UniProt accessions from the species-specific UniProt ID-mapping table, and each protein group was matched to a transcript when their UniProt accessions and/or curated gene symbols overlapped, with each match assigned a confidence tier (symbol-and-UniProt, UniProt-only, or symbol-only). All subsequent multi-omic analyses were performed on this combined MuData object. Because the gene-protein-group mapping is in some cases not definitive (179 gene symbols match more than one protein group and 323 protein groups match more than one gene symbol) the 4,166 matched gene symbols correspond to 4,350 gene–protein-group pairs. Compared gene-protein pairs vary therefore: per-pair for the correlation and variability comparisons (4,350 pairs, or 4,165 after requiring a finite within-condition CV²; 1,742 when both members are additionally required to be detected in ≥80 % of cells), and per-gene where protein groups are averaged within a gene (3,674 genes with usable pseudotime bins). The 3,627 genes reported as identified in both modalities (Figure S4K) apply additionally a detection filter of ≥10 % of a state’s cells in both modalities.

Multi-omic analyses: Four correlation metrics were used throughout, which are defined schematically in Figure S3A. mRNA-protein coupling was quantified per gene as the Pearson and Spearman correlation of the paired expression vectors across cells (per-gene correlation) or per cell across gene (per-cell correlation) (see Figure S3A), with p-values corrected for multiple testing by the Benjamini-Hochberg procedure over all tested feature pairs. Matched pairs were additionally ranked by their pooled per-gene correlation and split into deciles; mean transcript and protein abundance were then compared across deciles, and gene-set enrichment along the same ranking was projected onto them to show where each set concentrates. Per-cell S- and G2/M-phase scores for cell-cycle pseudotime were computed on the single-cell protein-abundance matrix using the canonical human Tirosh S-phase and G2/M-phase marker sets (43 and 54 genes, respectively; restricted to proteins quantified in ≥50 % of cells and log₂- transformed)^36^. A phase was scored only when ≥3 of its markers were quantified in a given cell. For each cell and phase, the score was computed with scanpy score_genes (v1.12.2) as the difference of the mean log₂ abundance of the phase markers and the mean of a background set of control proteins sampled from the same abundance bins (25 bins), so that scores are relative to each cell’s overall proteome rather than absolute. The discrete phase label was assigned as the higher-scoring phase, or G1 when both were ≤ 0. A continuous cell-cycle pseudotime was defined as the angular coordinate of each cell in the two-dimensional (S-score, G2/M-score) plane: θ = atan2(G2M − ⟨G2M⟩, S − ⟨S⟩), where ⟨·⟩ denotes the mean over paired cells, wrapped to [0, 2π). The angular origin was rotated so that the median angle of G1- labeled cells lay at 0, yielding a G1 → S → G2/M → G1 progression; cells were then ordered by θ for the gene-level trajectory plots. This pseudotime is therefore a cyclic, protein-derived coordinate in radians and is not a graph- or diffusion-based pseudotime. Cells whose S- or G2/M-score deviated by more than 4 standard deviations from the mean were excluded as scoring outliers (1 of 318 cells in the HeLa dataset).

Cell-to-cell variability was quantified per feature within each differentiation state as the coefficient of variation (CV = SD/mean) and its square (CV²), computed on measured values only, that is the unimputed protein layer, and requiring at least 15 detected cells per state for a feature to be scored. Matched gene– protein pairs were compared as the per-pair log₁₀ ratio of CV², taking each pair’s median across the states in which both members qualified. Local variability for the embedding overlays was computed per cell as the mean CV across that cell’s 30 nearest neighbors in protein PCA space, again on unimputed layers, either over all features of a modality or over a defined marker set.

Features changing between successive states were identified per modality by Wilcoxon rank-sum test with Benjamini–Hochberg correction, calling a feature changed at |log₂ fold change| > 0.5 and adjusted p < 0.05, and requiring at least 10 cells per group. Over-representation analysis used g:Profiler with Benjamini–Hochberg correction and term sizes restricted to 5–500 genes; gene-set enrichment analysis used GSEApy ‘prerank’ against GO:BP and GO:CC, reporting sets at FDR < 0.25. Cell diameter recorded by the dispenser at sorting was related to per-cell depth by Spearman correlation and used to bin cells by size at fixed cut points (13–18, 18–22 and 22–28 µm, equal spans and therefore unequal counts). Size-resolved enrichment combined two tests: per-bin one-versus-rest Wilcoxon up-features (FDR < 0.10, query capped at 300 genes) submitted to g:Profiler over-representation analysis against GO:BP and GO:CC with the whole identified transcriptome or proteome of that modality as background, and a binning-free gene-set enrichment analysis of all features ranked by Spearman correlation between expression and diameter, with positive scores indicating large-cell and negative small-cell enrichment. Transcripts detected in fewer than 25 cells were dropped beforehand; the protein layer was the imputed matrix.

Per-gene nascent signal was defined as the unspliced fraction Σunspliced/Σ(spliced+unspliced), scored only for genes with at least 50 total counts and detection in at least 20 cells; genes were then grouped into quartiles of that fraction and their mRNA–protein correlations compared across quartiles. Separately, to classify the shape of each matched pair along the trajectory, cells were divided into five equal-count pseudotime bins and each modality’s binned mean profile assigned a step pattern, yielding classes from synchronous through single-modality to offset and inverted. Program activity was scored per cell and modality with scanpy ‘score_genes’ over curated and Gene-Ontology-derived gene sets and plotted against pseudotime as a rolling mean ± s.e.m. Genes whose modalities diverged along pseudotime were selected as |Δρ| ≥ 0.40 between the RNA and protein Spearman correlations with pseudotime, with at least one modality significant at FDR < 0.05, and their mouse STRING interactions (combined score ≥ 400) clustered by the Louvain method (resolution 1.2), keeping components of at least three nodes and naming communities from CORUM complexes, Gene Ontology cellular components or hub genes.

A multi-omics factor analysis model (MOFA+, mofapy2 v0.7.4)^39^ was fitted at K = 15 latent factors, with factors contributing less than 1% variance in both modalities pruned before refitting. The imputed protein layer was not used, as it yields an additional factor loading on a small set of highly abundant proteins with no cross-modal concordance. Differentiation trajectories were reconstructed by diffusion pseudotime^37^ (scanpy, v1.12.2.) rooted independently in each modality at the naive cell whose PCA coordinates lie closest to the naive centroid. RNA velocity was estimated with scVelo [version 0.3.4]^41^ using the dynamical model, from UMI-deduplicated spliced and unspliced counts quantified with velocyto. Velocity speed was defined as the L2 norm of the per-cell velocity vector (|velocity|) and projected onto the protein UMAP embedding. Functional interpretation of gene and protein lists used over-representation analysis and gene-set enrichment analysis (GSEApy v1.3.0) against GO, KEGG, Reactome and WikiPathways gene sets, as well as g:Profiler (gprofiler-official v1.0.0).

Percentage of each modality’s full-sample component count that is recovered from cell subsets, as a function of the number of paired cells sampled: the number of principal components explaining 30% of that modality’s variance at n cells, expressed as a percentage of the value at all 495 matched cells (mean of 20 subsamples; 1,742 matched gene–protein pairs with both members detected in ≥80% of cells). Features were z-scored across cells and the component count taken as the interpolated crossing of the cumulative-variance curve, giving a fractional value. Cells were subsampled without replacement at each grid point with 20 replicates, sharing draws across modalities. Positional uncertainty along pseudotime was estimated with a Bayesian Gaussian-process latent-variable model (GPflow Bayesian GPLVM, one-dimensional latent, squared-exponential kernel, 50 inducing points)^70^ on the top 100 highly variable features per modality, z-scored per feature, with a per-cell Gaussian prior placing each state at equal spacing along the latent axis; the per-cell posterior standard deviation is reported. Fits used 3,000 iterations per modality (gradient tolerance 1e-6); the fitted observation-noise variance was 0.48 for RNA and 0.38 for protein (51.9% and 62.3% of variance explained) and the kernel length-scale 0.170 and 0.042.

For each functional sub-state program, coherence was defined as the mean pairwise correlation among that program’s own features across cells, computed within each condition, with every feature first residualized on per-cell depth (genes detected for RNA, features detected for protein). Protein used the unimputed layer with NaN-aware masked pairs, requiring at least 25 cells in which both members of a pair were detected. Significance was assessed against 200 size-matched random sets of matched pairs per program and modality. Features were excluded before scoring if flagged as common contaminants, or if they lay in the top 5% of proteome abundance while being transcribed in fewer than 25% of cells, and for the signaling pathways the ubiquitin–proteasome and ribosomal subunits shared between pathway supersets were likewise removed.

An independent single-cell proteomics experiment covering the naive, formative and primed stages (884 cells, sorted with the cellenONE, no paired transcriptome) was searched together with the multimodal proteomes in a single DIA-NN run, so that protein-group inference is shared between the datasets. Cells passed the same per-cell outlier filter as above. For the joint embedding, protein groups detected in at least 50 % of the pooled cells and in at least 10 % of each dataset’s cells were retained (3,166 groups), missing values were imputed (kNN) within each dataset separately so that no imputed value draws on the other experiment; each feature was standardized within each dataset, and the residual batch effect between experiments was removed with Harmony (harmonypy v0.2.0) using experiment as the covariate, after which a k-nearest-neighbor graph and UMAP were computed on the corrected components. Differential expression for the cross-dataset comparison was computed independently within each dataset as a per-protein Welch t-test between states on the measured log2 layer, Benjamini–Hochberg corrected per dataset and contrast; a protein counted as significant at adjusted p < 0.05 and |log2 fold change| > 0.5, and co-directionality is the fraction of proteins significant in both that changed in the same direction.

### Quantification and statistical analysis

All quantification and statistical analysis were performed in Python 3.12 using scanpy 1.12.2, anndata 0.12.18, mudata 0.3.9, muon 0.1.7, mofapy2 0.7.4, harmonypy 0.2.0, GSEApy 1.3.0, scikit-learn 1.9.0, SciPy 1.16.3, NumPy 2.4.6, pandas 2.3.3 and additional package versions listed above. All analysis parameters are specified in versioned YAML configuration files; the complete pipeline, configuration files and environment specification are available at GitHub (see *Code availability*). Unless stated otherwise, group comparisons used two-sided non-parametric tests (Wilcoxon rank-sum for unpaired and Wilcoxon signed-rank for paired comparisons), effect sizes are reported as medians with interquartile range, and multiple testing was controlled by the Benjamini–Hochberg procedure within each family of tests unless otherwise noted, and corrected values are reported as FDR. Correlations are Pearson or Spearman as stated. Resampling analyses used 20 replicates per grid point and permutation nulls 200 draws, with a fixed random seed (42) throughout. Box plots show the median and interquartile range (IQR) with whiskers extending to 1.5× IQR, and violin plots are clipped at the percentile stated in the corresponding legend. Sample sizes (n) denote the number of cells or the number of matched gene– protein pairs, as specified in each figure legend.

## Supplemental information

**Figure S1.**
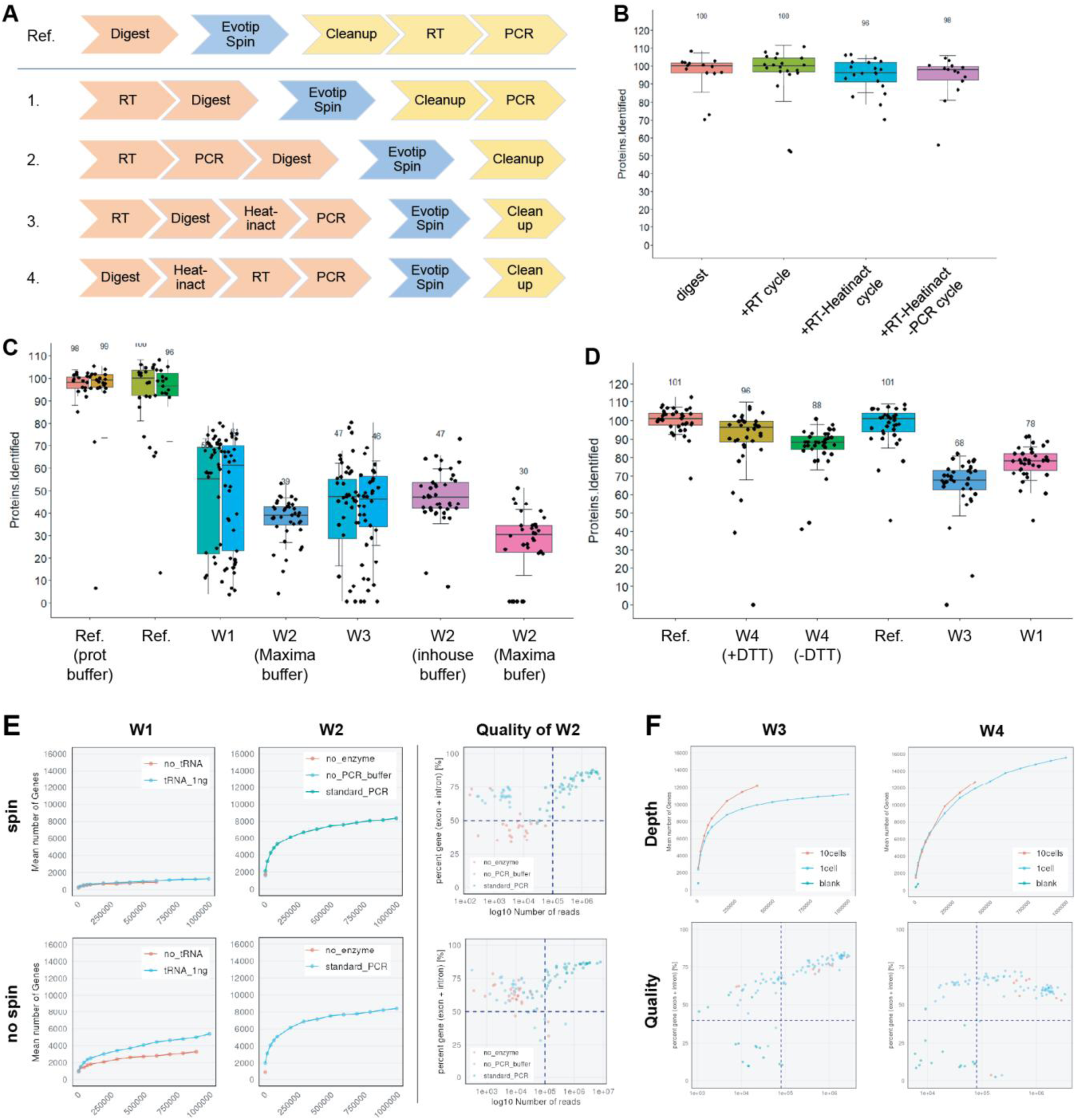
Systematic benchmarking of workflow step order and thermal conditions identifies an arrangement that preserves both proteome and transcriptome recovery. (A) Schematic of the candidate step orders evaluated. Four alternative arrangements (W1-4) that reposition reverse transcription (RT), pre-amplification (PCR), protein digestion, C18 capture and optional heat inactivation and cleanup relative to one another. A digest-only before C18 capture reference (protein digestion, C18 capture and following cleanup, RT, PCR) is shown on top. (B) Proteomic robustness to the thermal steps of the RNA workflow. Protein groups identified, normalized to a digest-only reference, after sequential addition of an RT thermal cycle, a heat-inactivation step, and a PCR thermal cycle. Median normalized identification remains between Sc and 100 across conditions, indicating that the added thermal steps do not substantially reduce proteome coverage. (C) Effect of step order on proteome recovery. Normalized protein groups identified across arrangements in which RT (W1) and PCR (W2 and W3) precede protein digestion and peptide capture, compared with digest-first references (Ref.). Digest-first conditions (Ref.) retain near-complete recovery (about 98-100%), whereas arrangements in which reverse transcription (W1 and W3) and, in particular, pre-amplification (W2) precede protein digest reduce recovery to about 30-47%. (D) Same as C in an independent experiment with testing reducing agents, confirming that digest-first orders outperform RT-first orders (digest-first arrangements: W4 with DTT 96% and without DTT 88%; RT-first arrangements: W3 68%, W1 78%) (E) Transcriptome recovery and library quality across arrangements, with (top) and without (bottom) C18-captured spin. Mean number of genes detected as a function of sequencing depth for the W1 (left) and W2 (center) arrangements, comparing carrier tRNA supplementation (no tRNA vs. tRNA_1ng). Library quality was checked as the fraction of gene reads (exon + intron) vs. read depth (right). (F) Same as E for the two lead arrangements, W3 (RT-digest-Heat-PCR, left) and W4 (Digest-Heat-RT-PCR, right).

**Figure S2.**
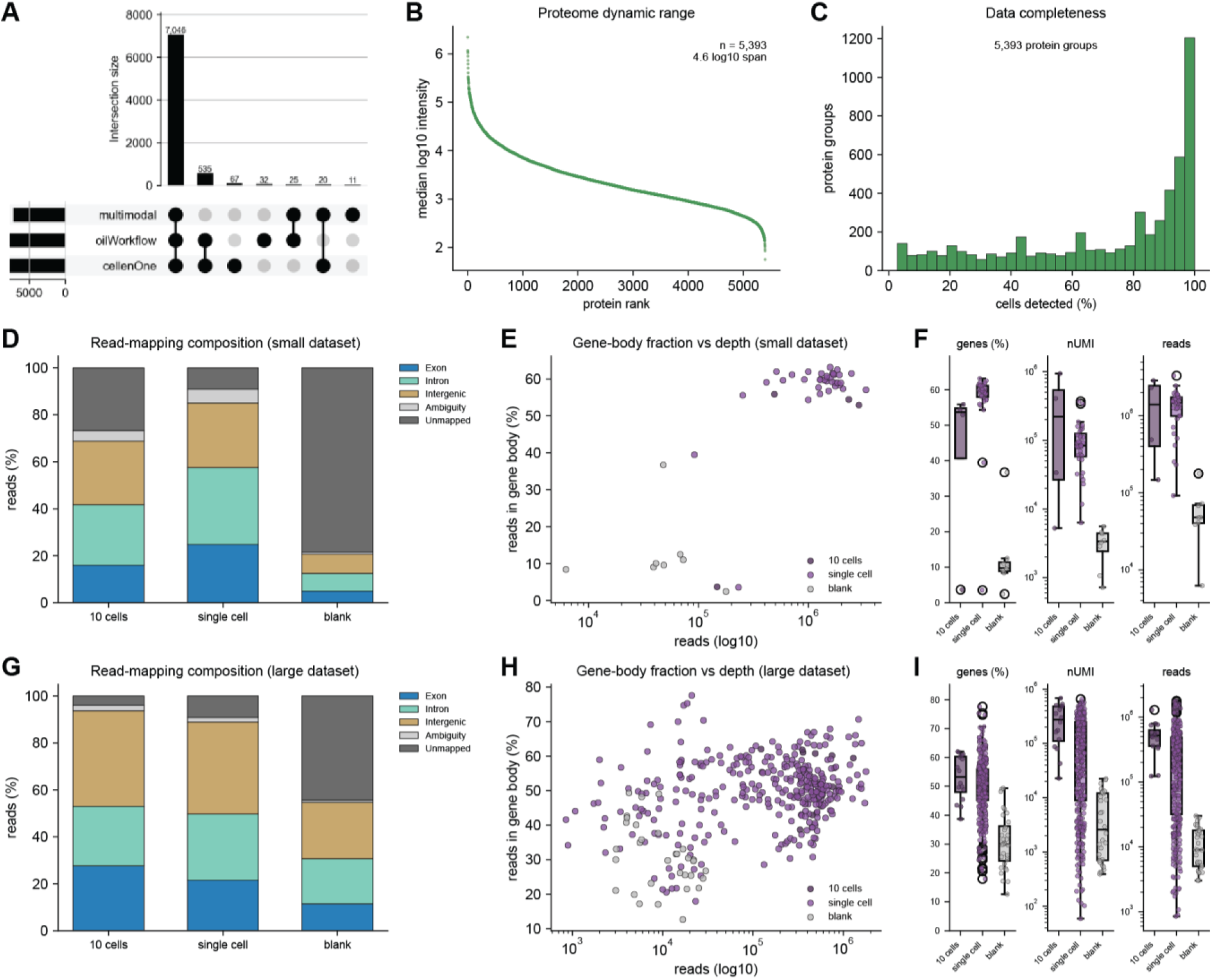
Single-cell proteomics and transcriptomics workflow characterization and quality control. (A) Comparison of single-cell proteomic identifications across all three workflows (cellenONE, oilWorkflow, multimodal). 7,046 protein groups are shared across all three workflows (91%). (B) Proteome dynamic range in the single-cell multimodal workflow (n = 5,393 protein groups spanning about 4.6 log10 units). (C) Data completeness: distribution of the fraction of single cells in which each protein group is detected; the peak near 100% corresponds to the core proteome detected in nearly every cell. (D-F) Quality control for transcriptomics on the small dataset (n = 72) across groups (10 cells / single cell / blank): (D) mean read-mapping composition (exon, intron, intergenic, ambiguous, and unmapped), (E) fraction of reads mapping to gene bodies (exon + intron) versus sequencing depth (reads, log10) per cell, and (F) reads-in-genes (%), unique molecular identifiers (nUMI) and total reads per cell. (G-I) Same as D-F for the large dataset (n = 318 cells, 3,904 overlapping genes). In all box plots the center line is the median, the box the interquartile range, and whiskers 1.5× IQR; individual cells or runs are overlaid as points.

**Figure S3.**
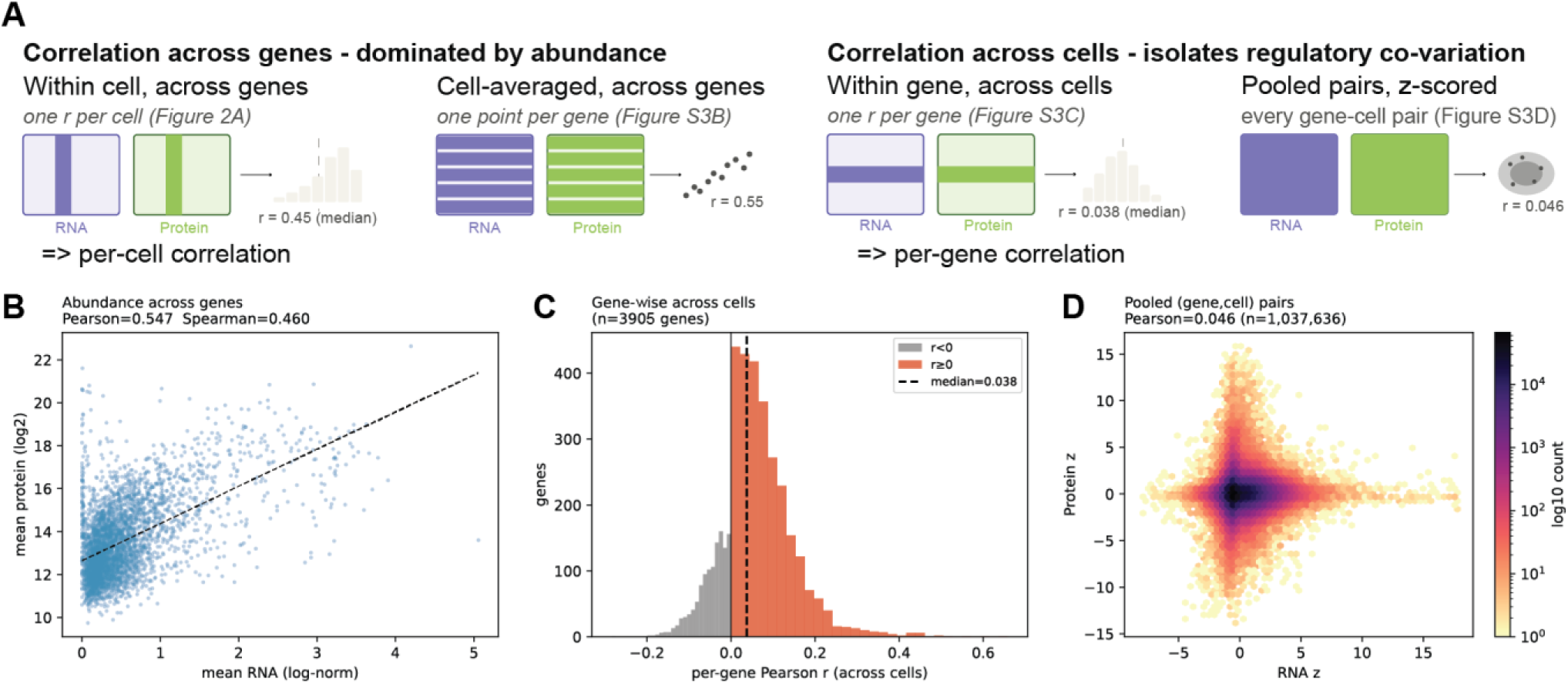
RNA-protein correlation metrics. (A) Schematic of the four correlation metrics used in this study. Rows are genes, columns are cells; highlighted entries indicate the values entering a single correlation. Metrics that correlate across genes (left) span the full between-gene abundance range and are therefore dominated by it, whether computed within one cell (per-cell correlation, Fig. 2A) or on cell-averaged values (B). Metrics that operate per-gene across cells (right) display regulatory co-variation, either within one gene (per-gene correlation, C) or by z-scoring on gene-cell pairs (D). (B) Abundance correlation across genes: each gene’s mean transcript level (log-normalized) versus its mean protein level (log₂) averaged within cells (each point is one gene; Pearson r = 0.55, Spearman ρ = 0.46); measures whether abundant mRNAs yield abundant proteins. (C) Per-gene correlation across cells: distribution of the per-gene Pearson r computed across cells (one value per gene; n = 3,904 genes; median r = 0.038 (dashed line); gray = r < 0, red = r ≥ 0). This metric is an order of magnitude weaker than the abundance correlation in (B). (D) Pooled (gene, cell) pair correlation: 2D density (log-scaled hexbin) distribution of all per-gene z-scored (transcript, protein) pairs per cell (n = 1,037,636 pairs; Pearson r = 0.046).

**Figure S4.**
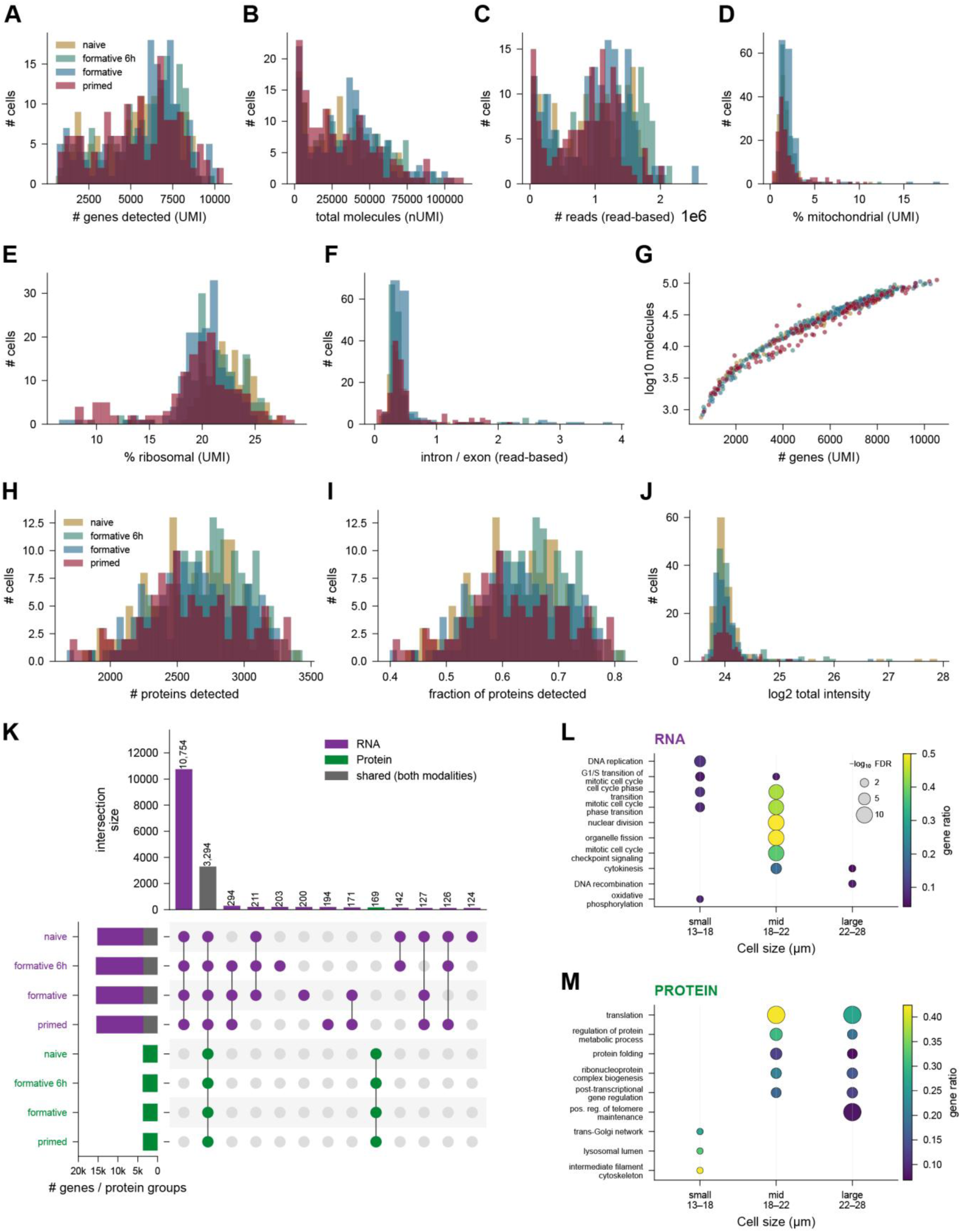
Quality control and differentiation validation. (A–G) Per-cell transcriptome quality control computed on the deduplicated UMI molecule-count matrix (n = 612 cells), colored by differentiation state: genes detected (A), total molecules (B), sequencing reads (C, read-based), percent mitochondrial (D) and ribosomal (E) UMIs, intron/exon read ratio (F, read-based), and per-cell coverage as genes versus total molecules (G). Recomputing depth on molecule counts derived from the whole transcriptome dataset (612 cells, including cells which are not in the proteomics dataset due to applied filters) rather than reads lowers the median genes per cell from ∼10,090 to ∼6,160 (UMI deduplication) while leaving percent mitochondrial essentially unchanged (1.65 versus 1.66). (H–J) Per-cell proteome quality control (n = 540 cells): protein groups detected (H), fraction of the quantified proteome detected (I), and log2 total sample intensity (J). (K) UpSet plot of feature identification across the four states and both modalities. RNA (restricted to protein-coding genes) and protein are collapsed to a common gene-symbol universe, and a feature is called identified in a state if detected in at least 10 % of that state’s cells. Left set-size bars (# genes / protein groups) split each RNA state set into the fraction also identified in the proteome (grey) and the RNA-only remainder (purple); protein sets are green. Intersection bars are RNA-only (purple), protein-only (green) or shared across modalities (grey); most features are shared across all four states within a modality, and 3,627 genes are identified in both the transcriptome and the proteome. (L, M) Cell-size-resolved functional enrichment for RNA (L) and protein (M). Pooling all four states, cells are binned by cellenONE diameter into 13–18 / 18–22 / 22–28 µm groups (equal size spans, so cell counts differ); displayed terms are drawn from two complementary tests shown in one dot plot, per-tertile GO over-representation (each term plotted in every size bin where it is significant) and continuous size-ranked GSEA (each term placed in a single bin by the sign of its enrichment along the size axis). Dot size = −log10 FDR (size legend in L), color = gene ratio.

**Figure S5.**
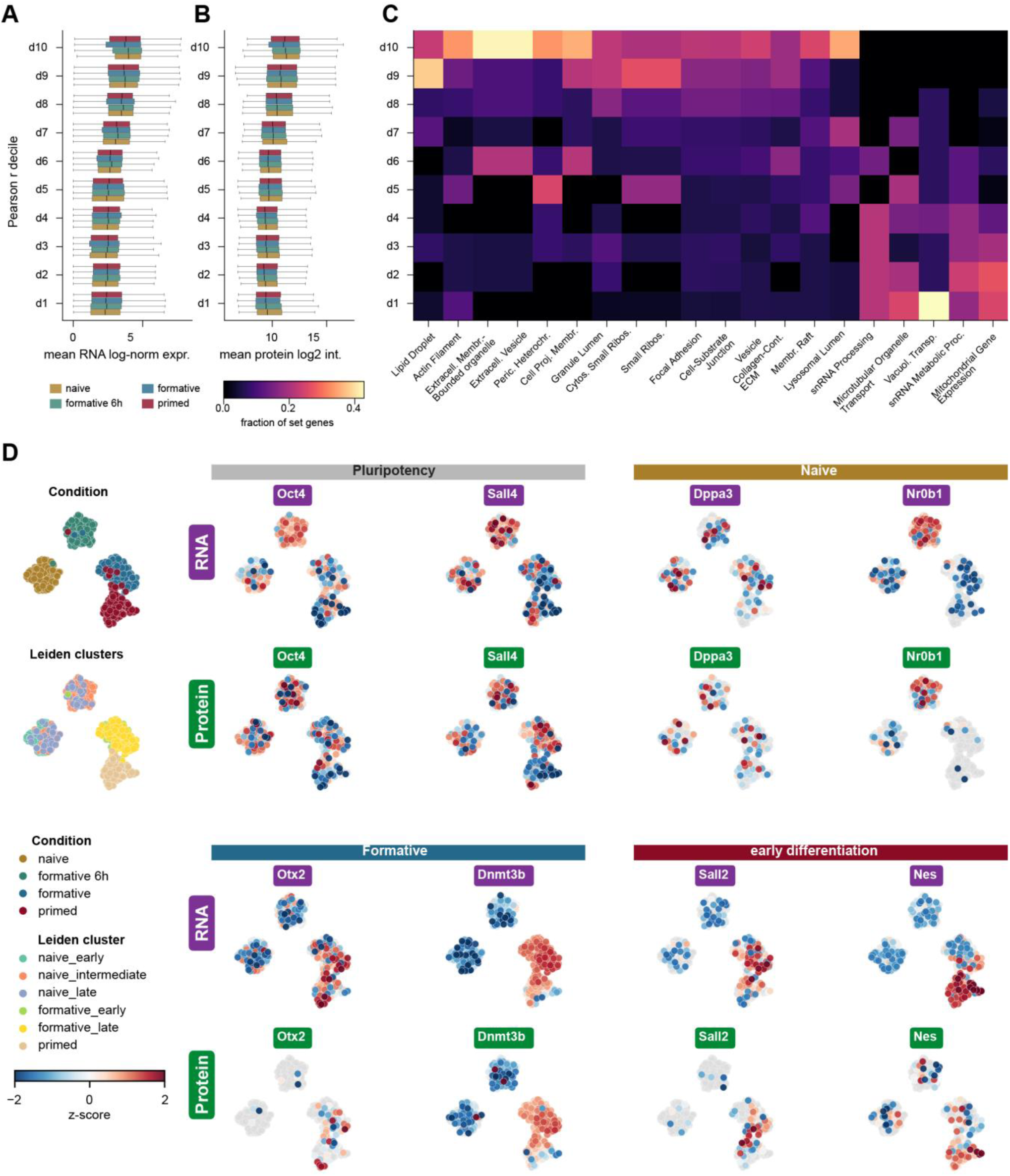
RNA-protein correlation gradient and RNA-based UMAP coordinates. (A, B) Gene–protein pairs were split into deciles by their pooled per-cell Pearson correlation (d1 = lowest r, d10 = highest r; the pooled r span is printed on the axis). For each decile, boxplots (median, IQR, 1.5×IQR whiskers) show the distribution of per-gene mean RNA log-normalized expression (A) and mean protein log2 intensity (B), split by state (shared legend, right of B); abundance rises monotonically from d1 to d10 in every state and both modalities, so the pairs whose RNA and protein co-vary across single cells are also the more abundant ones. (C) Functional programs along the pooled correlation gradient. GSEA of the full pooled r ranking identifies the significantly enriched sets; for each set the heatmap projects the fraction of its member genes falling in each r-decile, showing where along the gradient it concentrates. Sets loaded at the highly-correlated, abundant tail (ribosomal subunits, actin filament, extracellular vesicle, lipid droplet) accumulate in d9–d10, whereas the weakly-correlated sets (mitochondrial gene expression, snRNA processing, vacuolar / organelle transport) accumulate in d1–d4. (D) Extended lineage-marker maps, all on the RNA UMAP so RNA and protein of each marker are directly comparable on one layout. Eight markers grouped by program, each with RNA (top of a block, purple chip) and protein (bottom, green chip) z-scored on the unimputed layers over detected cells only (undetected cells grey); the reference column shows the same RNA UMAP colored by state (top) and by the six protein Leiden sub-states (bottom). The protein-derived sub-states and states map coherently onto the RNA embedding; pluripotency factors (Oct4, Sall4) are high in the naive cluster and decline while Dnmt3b and Otx2 rise toward the primed cluster, concordantly in RNA and protein where protein is detected. Several markers are protein-sparse (Otx2 14 %, Sall2 17 %, Dppa3 23 %, Nr0b1 30 %, Nes 35 % of cells versus RNA 44–100 %), reflecting the lower per-cell depth of single-cell proteomics rather than absence of the transcript.

**Figure S6.**
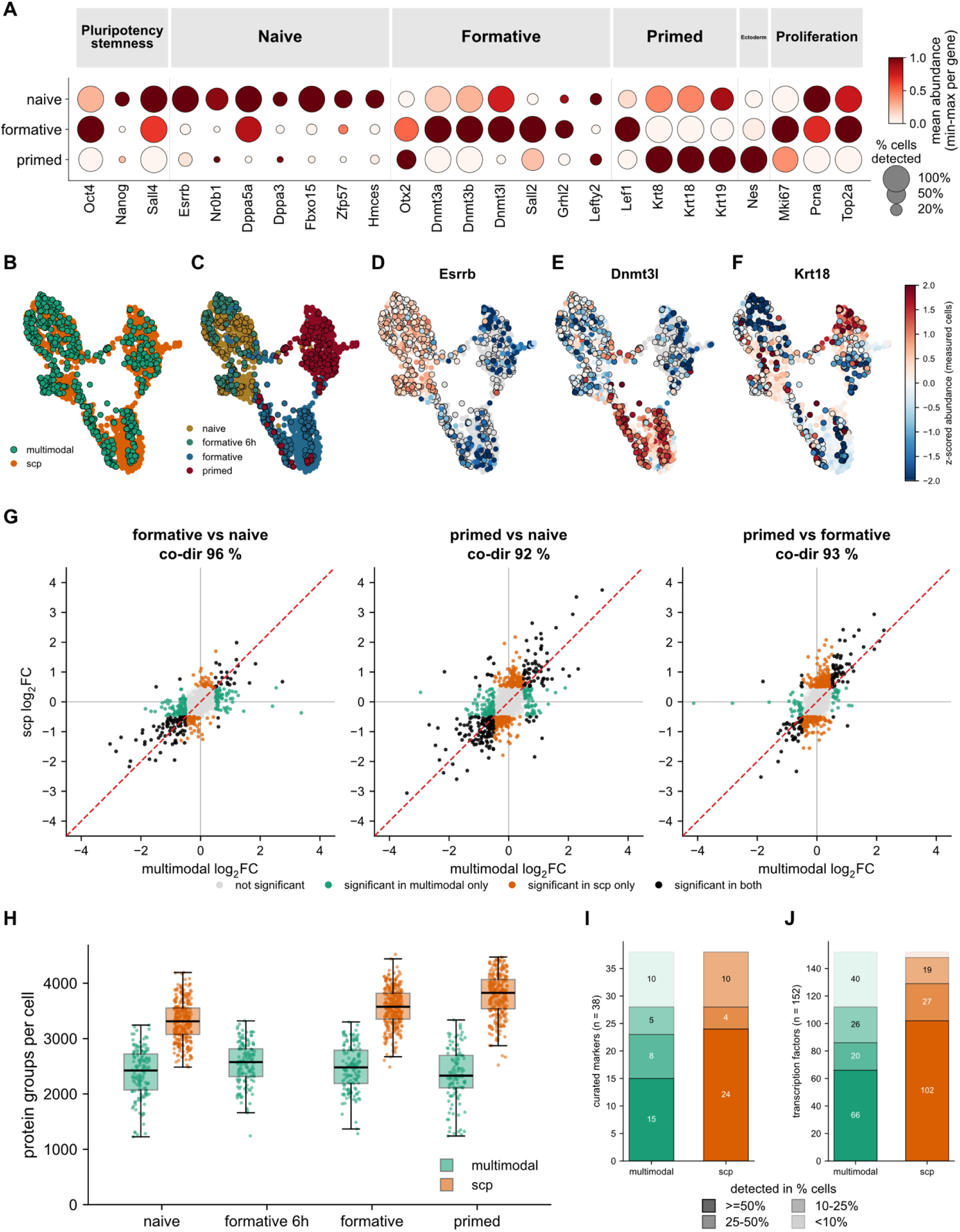
Single-cell proteomics (scp) integrates into and reproduces the multimodal proteome differentiation program in independent mESC differentiation dataset (884 cells: naive 298, formative 313, primed 273). (A) Marker dot-plot for the scp dataset across its three states. Only markers detected in the multimodal proteome are shown (see Figure 3H); dot color is the per-gene min–max-normalized mean of the log2 (non-imputed) intensity, and dot size is its detection in percentage of cells. Markers are grouped by state category (pluripotency/stemness, naive, formative, primed, ectoderm, proliferation). (B-F) Joint UMAP of both datasets, computed after retaining protein groups detected in ≥50% of the pooled cells and ≥10% of each dataset’s cells, imputing within each dataset separately, standardizing every feature within each dataset, and removing the residual between-experiment batch effect with Harmony. Colored by dataset (B) and by differentiation state (C): the two experiments overlap within every state rather than separating by experiment, and the states remain ordered from naive through formative to primed. (D–F) Measured abundance of Esrrb (D), Dnmt3l (E) and Krt18 (F) on the same coordinates, z-scored per protein across both datasets over the cells in which it was measured; cells in which it was not detected are gray. (G) Cross-dataset concordance of differential expression, computed independently within each dataset: per-protein Welch t-test between states on the 2,240 protein groups detected in ≥50% of the cells of both datasets, for the three shared contrasts (formative vs naive, left; primed vs naive, middle; primed vs formative, right). Each point is a protein group (x = multimodal log2FC, y = scp log2FC). Color gives the significance call in each dataset (Benjamini–Hochberg adjusted p < 0.05 **and** |log2FC| > 0.5): green = multimodal only, orange = scp only, black = both, gray = neither. The percentage under each contrast is the fraction of the proteins significant in both datasets that change in the same direction (co-dir). (H) Protein groups quantified per cell by state and dataset (box = median and IQR, whiskers = 1.5×IQR, points = individual cells), recomputed on the shared protein-group space. The scp dataset is deeper in every shared state (Mann–Whitney p ≤ 5×10⁻³¹). The 6 h formative intermediate exists only in the multimodal series. (I, J) Detection completeness of curated lineage markers (I) and annotated transcription factors (J) in each dataset. Bar height is the number of list members present in the joint search space (37 and 146, respectively), partitioned by the fraction of cells in which each was detected. Raw identification counts do not separate the two workflows since both quantify essentially the same features (36 versus 37 markers, 145 versus 146 transcription factors), whereas completeness does.

