## Supplemental Figures for "In the same cell, the proteome defines cellular state and the transcriptome marks transitions"

### Supplemental information

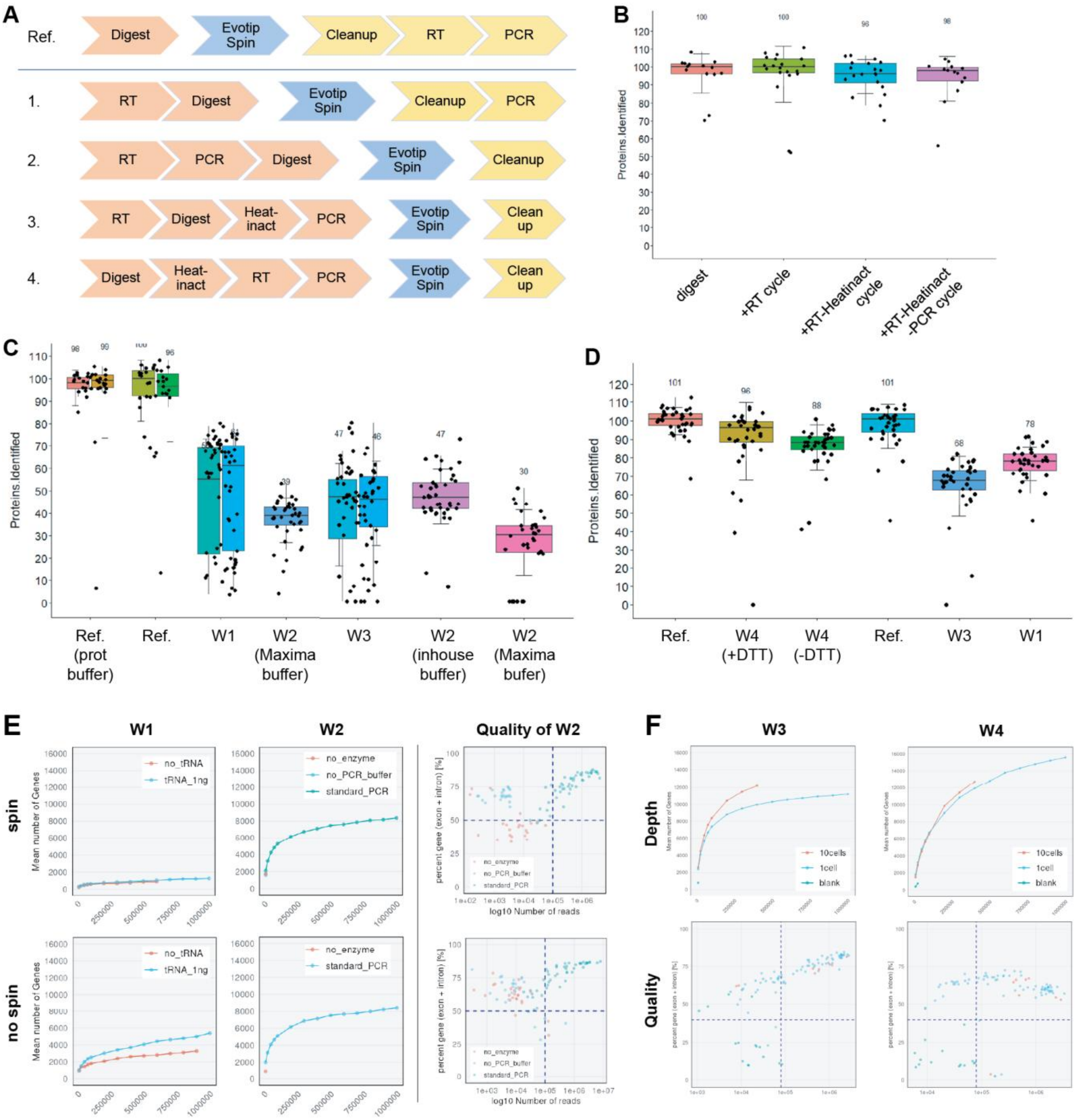


***Figure S1 | Systematic benchmarking of workflow step order and thermal conditions identifies an arrangement that preserves both proteome and transcriptome recovery.*** *(A) Schematic of the candidate step orders evaluated. Four alternative arrangements (W1-4) that reposition reverse transcription (RT), pre-amplification (PCR), protein digestion, C18 capture and optional heat inactivation and cleanup relative to one another. A digest-only before C18 capture reference (protein digestion, C18 capture and following cleanup, RT, PCR) is shown on top. (B) Proteomic robustness to the thermal steps of the RNA workflow. Protein groups identified, normalized to a digest-only reference, after sequential addition of an RT thermal cycle, a heat-inactivation step, and a PCR thermal cycle. Median normalized identification remains between 96 and 100 across conditions, indicating that the added thermal steps do not substantially reduce proteome coverage. (C) Effect of step order on proteome recovery. Normalized protein groups identified across arrangements in which RT (W1) and PCR (W2 and W3) precede protein digestion and peptide capture, compared with digest-first references (Ref.). Digest-first conditions (Ref.) retain near-*

*complete recovery (about 98-100%), whereas arrangements in which reverse transcription (W1 and W3) and, in particular, pre-amplification (W2) precede protein digest reduce recovery to about 30-47%. (D) Same as C in an independent experiment with testing reducing agents, confirming that digest-first orders outperform RT-first orders (digest-first arrangements: W4 with DTT 96% and without DTT 88%; RT-first arrangements: W3 68%, W1 78%) (E) Transcriptome recovery and library quality across arrangements, with (top) and without (bottom) C18-captured spin. Mean number of genes detected as a function of sequencing depth for the W1 (left) and W2 (center) arrangements, comparing carrier tRNA supplementation (no tRNA vs. tRNA_1ng). Library quality was checked as the fraction of gene reads (exon + intron) vs. read depth (right). (F) Same as E for the two lead arrangements, W3 (RT-digest-Heat-PCR, left) and W4 (Digest-Heat-RT-PCR, right).*

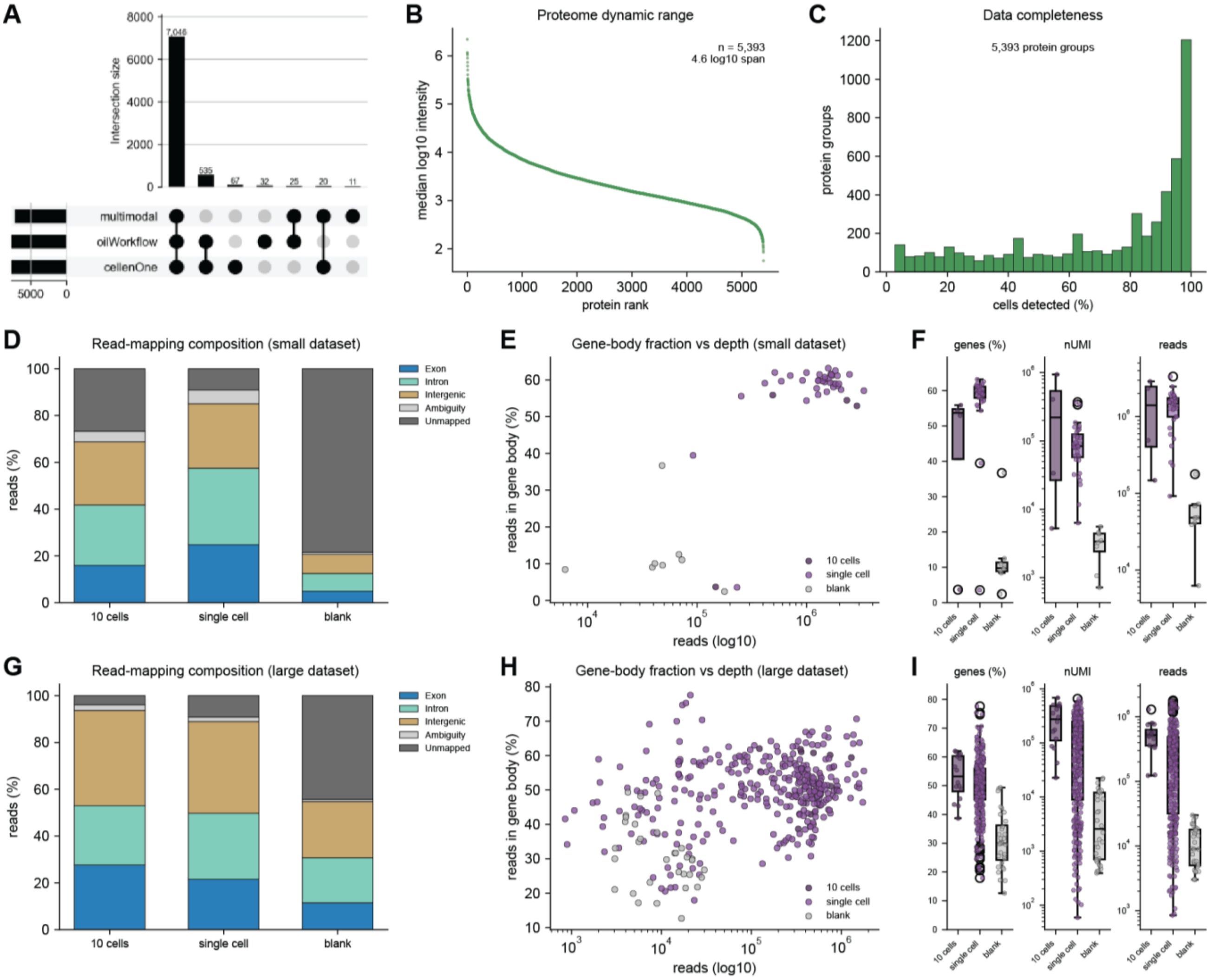


**Figure S2 | Single-cell proteomics and transcriptomics workflow characterization and quality control.** *(A) Comparison of single-cell proteomic identifications across all three workflows (cellenONE, oilWorkflow, multimodal). 7,046 protein groups are shared across all three workflows (91%). (B) Proteome dynamic range in the single-cell multimodal workflow (n = 5,393 protein groups spanning about 4.6 log10 units). (C) Data completeness: distribution of the fraction of single cells in which each protein group is detected; the peak near 100% corresponds to the core proteome detected in nearly every cell. (D-F) Quality control for transcriptomics on the small dataset (n = 72) across groups (10 cells / single cell / blank): (D) mean read-mapping composition (exon, intron, intergenic, ambiguous, and unmapped), (E) fraction of reads mapping to gene bodies (exon + intron) versus sequencing depth (reads, log10) per cell, and (F) reads-in-genes (%), unique molecular identifiers (nUMI) and total reads per cell. (G-I) Same as D-F for the large dataset (n = 318 cells, 3,904 overlapping genes). In all box plots the center line is the median, box the interquartile range, and whiskers 1.5× IQR; individual cells or runs are overlaid as points.*

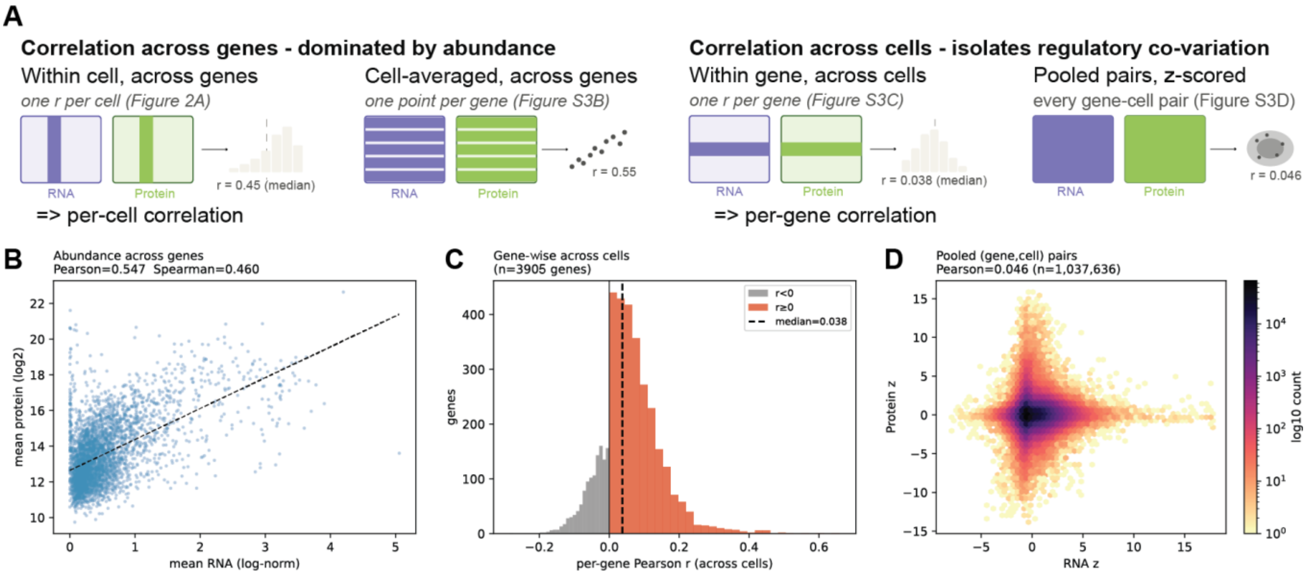


***Figure S3 | RNA-protein correlation metrics.*** *(A) Schematic of the four correlation metrics used in this study. Rows are genes, columns are cells; highlighted entries indicate the values entering a single correlation. Metrics that correlate across genes (left) span the full between-gene abundance range and are therefore dominated by it, whether computed within one cell (per-cell correlation, Fig. 2A) or on cell-averaged values (B). Metrics that operate per-gene across cells (right) display regulatory co-variation, either within one gene (per-gene correlation, C) or by z-scoring on gene-cell pairs (D). (B) Abundance correlation across genes: each gene's mean transcript level (log-normalized) versus its mean protein level ($log_2$) averaged within cells (each point is one gene; Pearson r = 0.55, Spearman ρ = 0.46); measures whether abundant mRNAs yield abundant proteins. (C) Per-gene correlation across cells: distribution of the per-gene Pearson r computed across cells (one value per gene; n = 3,904 genes; median r = 0.038 (dashed line); gray = r < 0, red = r ≥ 0). This metric is an order of magnitude weaker than the abundance correlation in (B). (D) Pooled (gene, cell) pair correlation: 2D density (log-scaled hexbin) distribution of all per-gene z-scored (transcript, protein) pairs per cell (n = 1,037,636 pairs; Pearson r = 0.046).*

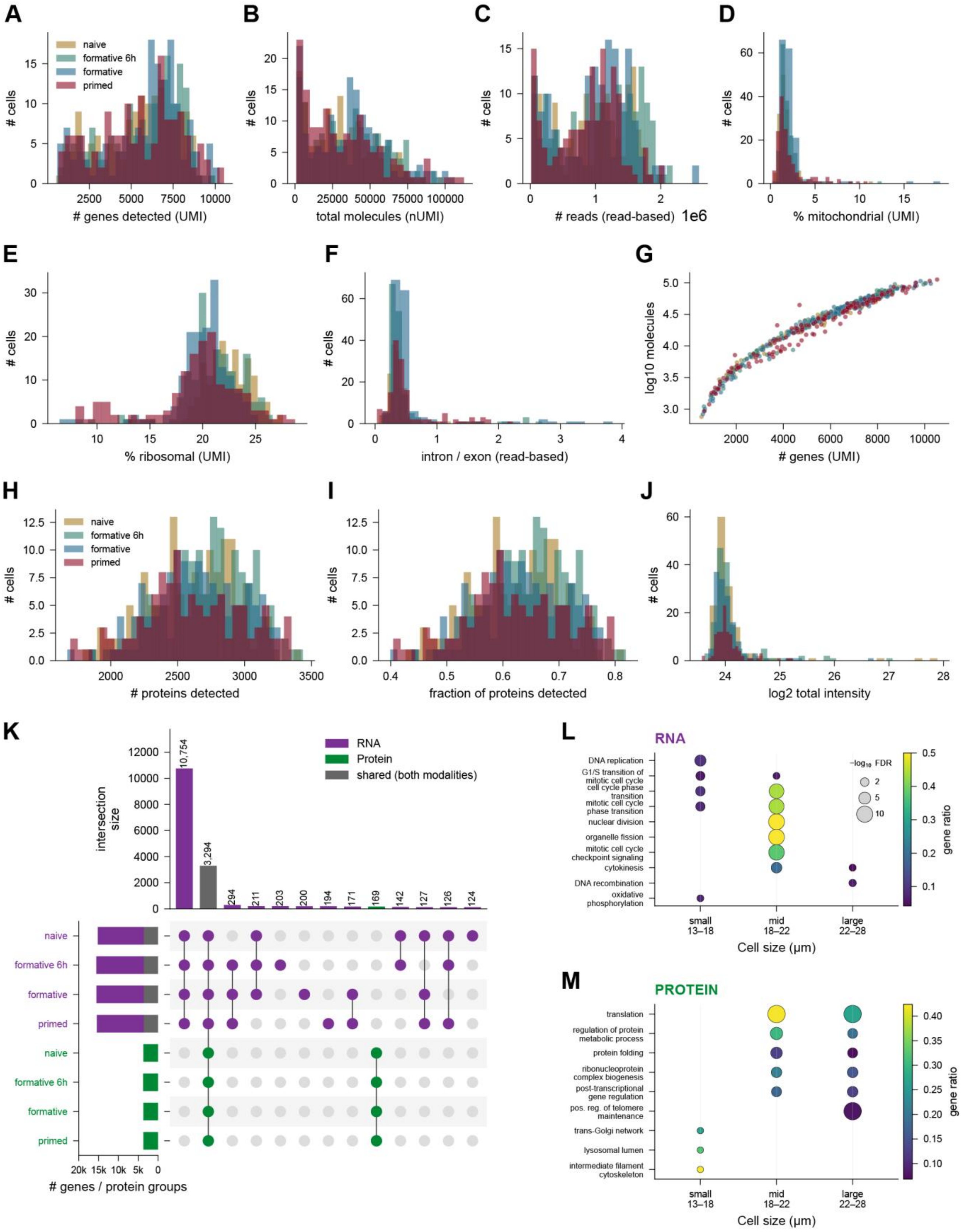


***Figure S4 | Quality control and differentiation validation.*** *(A–G) Per-cell transcriptome quality control computed on the deduplicated UMI molecule-count matrix (n = 612 cells), colored by differentiation state: genes detected (A), total molecules (B), sequencing reads (C, read-based), percent mitochondrial (D) and ribosomal (E) UMIs, intron/exon read ratio (F, read-based), and per-cell coverage as genes versus total molecules (G). Recomputing depth on molecule counts derived from the whole*

*transcriptome dataset (612 cells, including cells which are not in the proteomics dataset due to applied filters) rather than reads lowers the median genes per cell from ~10,090 to ~6,160 (UMI deduplication) while leaving percent mitochondrial essentially unchanged (1.65 versus 1.66). (H–J) Per-cell proteome quality control (n = 540 cells): protein groups detected (H), fraction of the quantified proteome detected (I), and log2 total sample intensity (J). (K) UpSet plot of feature identification across the four states and both modalities. RNA (restricted to protein-coding genes) and protein are collapsed to a common gene-symbol universe, and a feature is called identified in a state if detected in at least 10 % of that state's cells. Left set-size bars (# genes / protein groups) split each RNA state set into the fraction also identified in the proteome (grey) and the RNA-only remainder (purple); protein sets are green. Intersection bars are RNA-only (purple), protein-only (green) or shared across modalities (grey); most features are shared across all four states within a modality, and 3,627 genes are identified in both the transcriptome and the proteome. (L, M) Cell-size-resolved functional enrichment for RNA (L) and protein (M). Pooling all four states, cells are binned by cellenONE diameter into 13–18 / 18–22 / 22–28 µm groups (equal size spans, so cell counts differ); displayed terms are drawn from two complementary tests shown in one dot plot, per-tertile GO over-representation (each term plotted in every size bin where it is significant) and continuous size-ranked GSEA (each term placed in a single bin by the sign of its enrichment along the size axis). Dot size = −log10 FDR (size legend in L), color = gene ratio.*

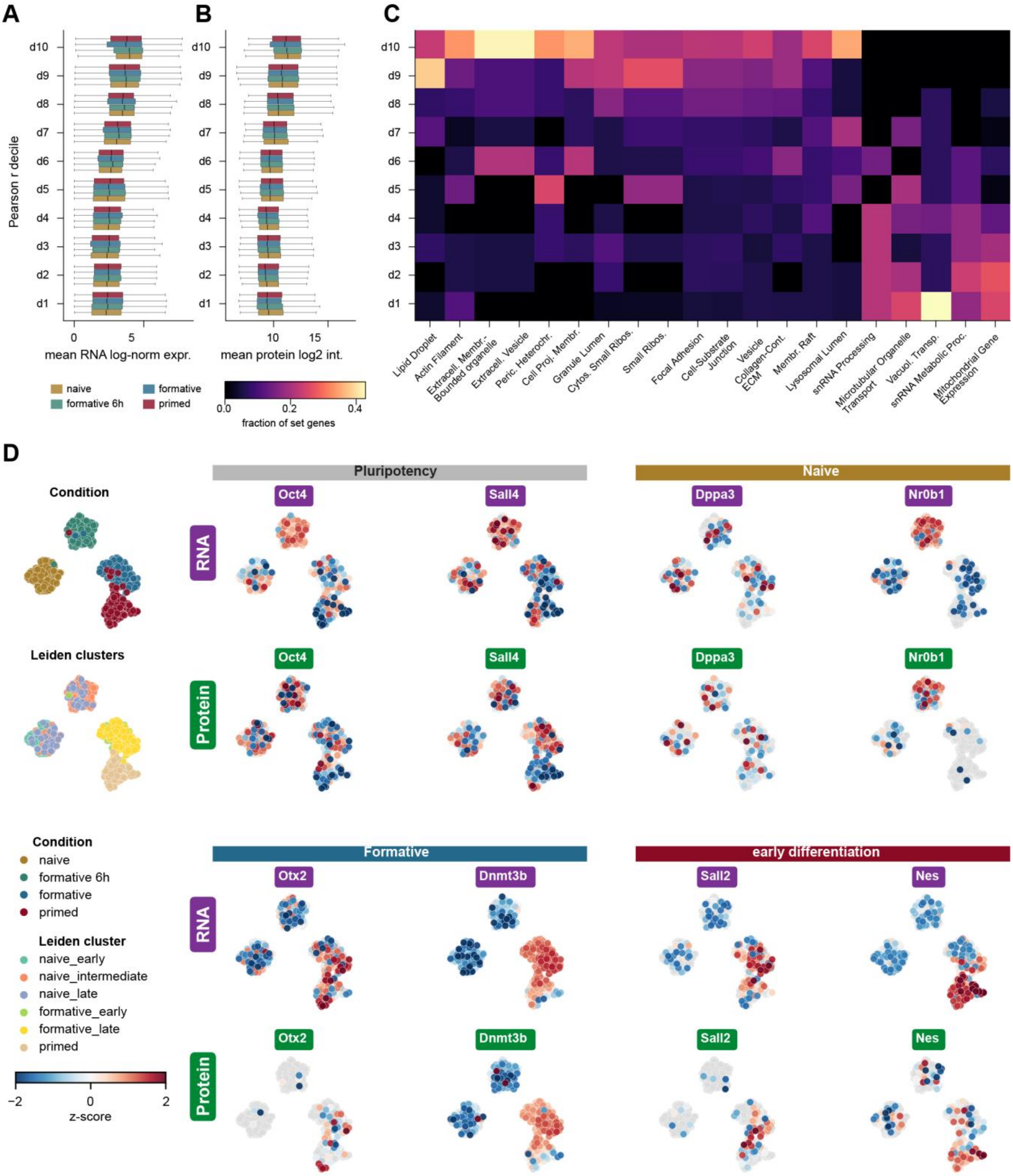


***Figure S5 | RNA-protein correlation gradient and RNA-based UMAP coordinates.*** *(A, B) Gene–protein pairs were split into deciles by their pooled per-cell Pearson correlation (d1 = lowest r, d10 = highest r; the pooled r span is printed on the axis). For each decile, boxplots (median, IQR, 1.5×IQR whiskers) show the distribution of per-gene mean RNA log-normalized expression (A) and mean protein log2 intensity (B), split by state (shared legend, right of B); abundance rises monotonically from d1 to d10 in every state and both modalities, so the pairs whose RNA and protein co-vary across single cells are also the more abundant ones. (C) Functional programs along the pooled correlation gradient. GSEA of the full pooled r ranking identifies the significantly enriched sets; for each set the heatmap projects the fraction of its member genes falling in each r-decile, showing where along the gradient it concentrates. Sets loaded at the highly-correlated, abundant tail (ribosomal subunits, actin filament, extracellular vesicle, lipid droplet) accumulate in d9–d10, whereas the weakly-correlated sets (mitochondrial gene expression, snRNA*

*processing, vacuolar / organelle transport) accumulate in d1–d4. (D) Extended lineage-marker maps, all on the RNA UMAP so RNA and protein of each marker are directly comparable on one layout. Eight markers grouped by program, each with RNA (top of a block, purple chip) and protein (bottom, green chip) z-scored on the unimputed layers over detected cells only (undetected cells grey); the reference column shows the same RNA UMAP colored by state (top) and by the six protein Leiden sub-states (bottom). The protein-derived sub-states and states map coherently onto the RNA embedding; pluripotency factors (Oct4, Sall4) are high in the naive cluster and decline while Dnmt3b and Otx2 rise toward the primed cluster, concordantly in RNA and protein where protein is detected. Several markers are protein-sparse (Otx2 14 %, Sall2 17 %, Dppa3 23 %, Nr0b1 30 %, Nes 35 % of cells versus RNA 44–100 %), reflecting the lower per-cell depth of single-cell proteomics rather than absence of the transcript.*

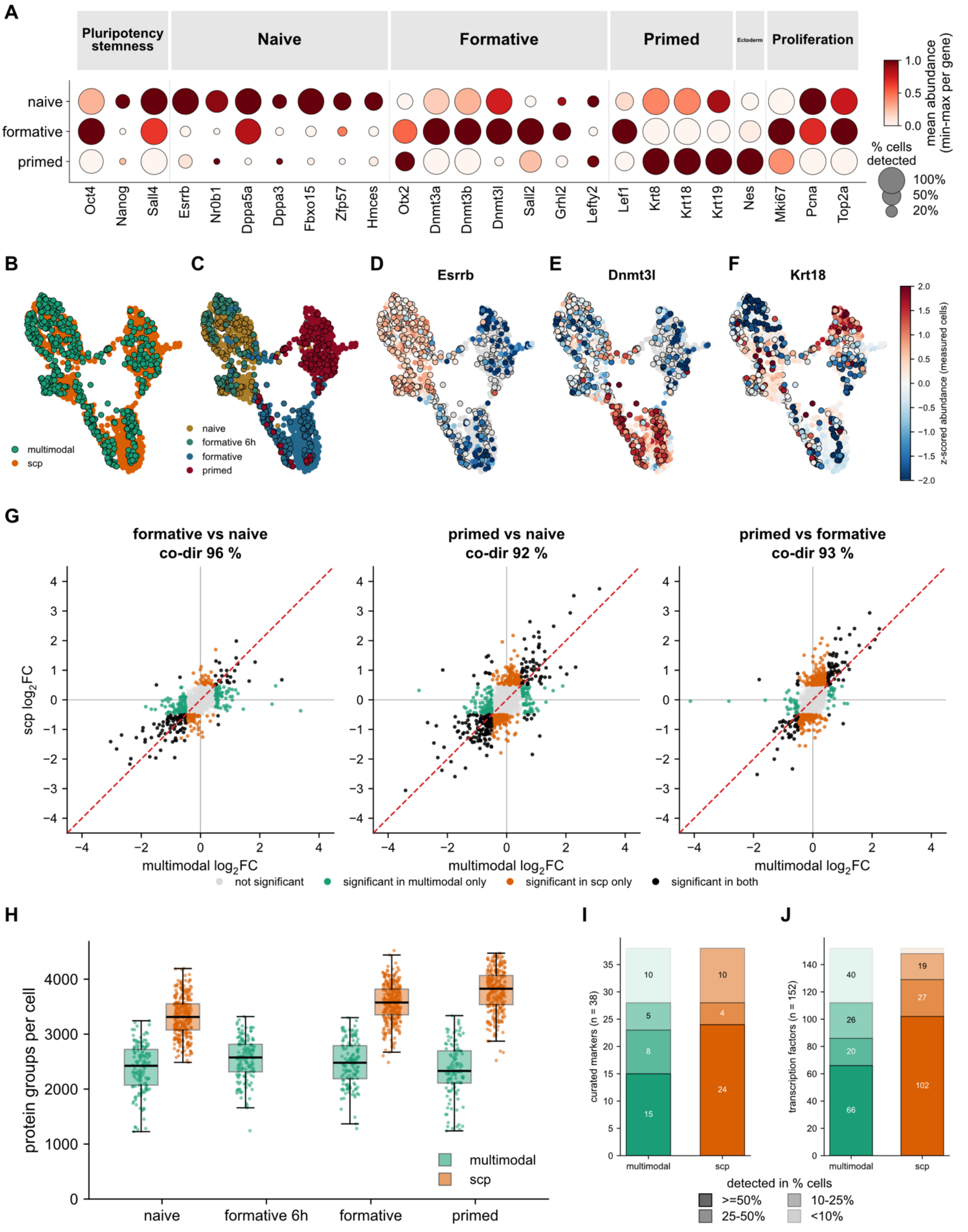


***Figure S6 | Single-cell proteomics (scp) integrates into and reproduces the multimodal proteome differentiation program in independent mESC differentiation dataset (884 cells: naive 298, formative 313, primed 273).*** *(A) Marker dot-plot for the scp dataset across its three states. Only markers detected in the multimodal proteome are shown (see Figure 3H); dot color is the per-gene min–max-normalized mean of the log2 (non-imputed) intensity, and dot size is its detection in percentage of cells. Markers are grouped by state category (pluripotency/stemness, naive, formative, primed, ectoderm, proliferation). (B-F) Joint*
